# Regulation of YAP activity by nuclear G-actin binding

**DOI:** 10.1101/2025.07.30.667612

**Authors:** Hong Wang, Irma M. Jayawardana, Justus M. Fleisch, Dennis Frank, Bing Zhao, Christos Kamaras, Peter Gebhardt, Julian Knerr, Shamphavi Sivabalasarma, Sonja-Verena Albers, Robert Grosse

## Abstract

Graphical Abstract:
YAP functions as a G-actin binding protein. Three high affinity actin binding sites, L65, L68 and W199 are identified in YAP that when mutated into YAP^DDY^ attenuate YAP/actin interaction as well as YAP activity. Thus, nuclear actin binding to YAP is necessary for its transcriptional activity as a co-activator of TEAD. Incorporation of G-actin into the YAP/TEAD complex is predicted to induce TEAD conformational change involving a 4α interface. Therefore, the formation of a dynamic TEAD/YAP/actin ternary complex necessary for transcription is proposed.

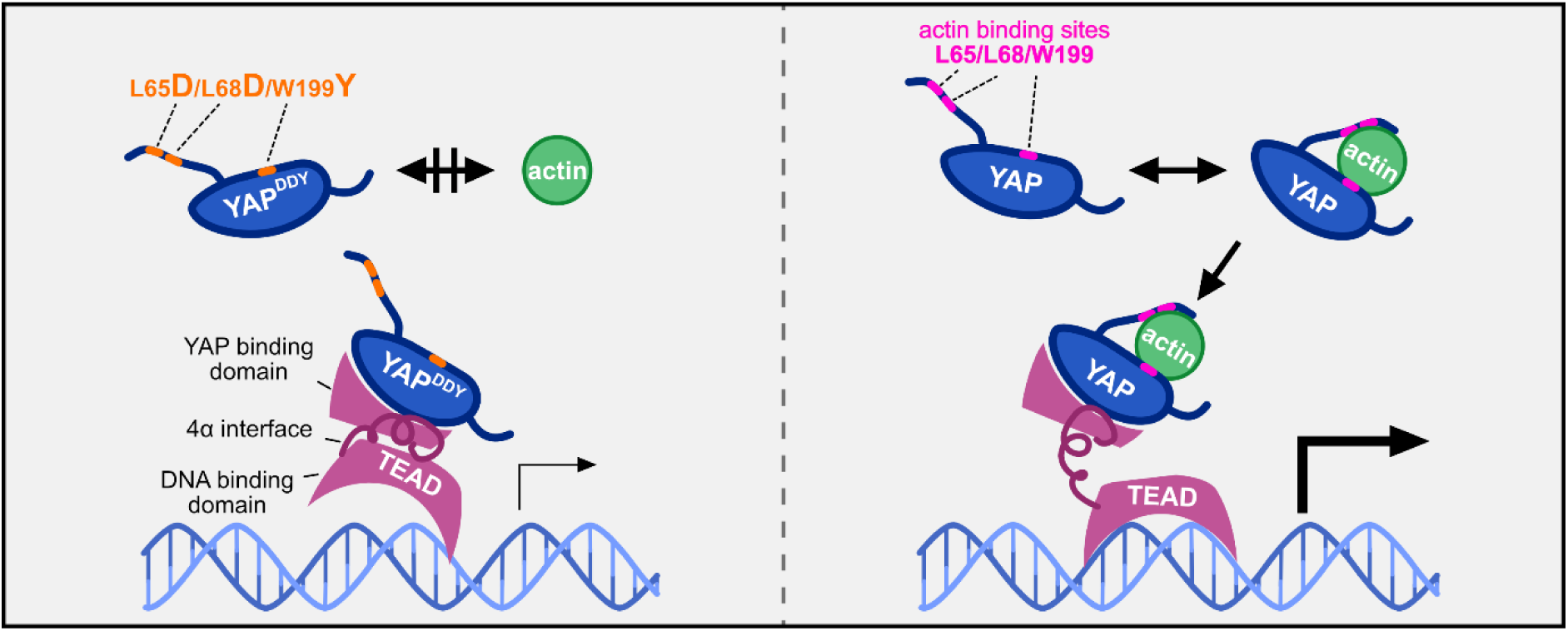

The Yes-associated protein YAP belongs to the TEAD (TEA/ATTS domain) transcriptional co-activators that shuttle between cytoplasm and nuclear compartment. YAP and its paralog TAZ (transcriptional co-activator with PDZ-binding motif) play essential roles in the Hippo pathway to control tissue and organ size. In addition, YAP is critically involved in numerous cellular processes such as differentiation, proliferation, cell migration and cancer metastasis as well as mechanotransduction and cytoskeletal dynamics. The actin cytoskeleton controls YAP activity in multiple ways via tensile forces, cell density and cell-cell adhesion as well as shear stress or other biomechanical cues. Here we discover YAP as a novel G-actin binding protein. We identify three high affinity YAP actin binding sites involving critical residues within the N-terminus of YAP. Moreover, actin binding to YAP is necessary for its transcriptional activity and function such as during cell density control. Mutation of the three actin binding residues results in a loss of nuclear YAP co-activator function towards TEAD while actin binding to YAP is required for TEAD target gene regulation. Our data point towards the formation of a dynamic TEAD/YAP/actin ternary complex necessary for transcription.

## INTRODUCTION

Dynamic rearrangement of nuclear actin has emerged as an integral component of the nuclear skeleton serving functions in chromatin dynamics, genome organization, DNA repair and transcription(1–6). Notably, many of these functions are due to actin turnover and hence the ability of actin to assemble and disassemble filaments (7). In addition, dynamic G-actin interactions have been shown to control specific transcription factors. One prominent example is the SRF activator MRTF (also known as MAL, MKL1) (8). MRTF interacts with G-actin, both in the cytoplasm and in the nucleus (9, 10). While dissociation of G-actin from the N-terminal RPEL domain from MRTF exposes its nuclear localization signal for nuclear import, G-actin loading inside the nucleus in turn is necessary for efficient MRTF export (11–14). Therefore, cytoskeleton dynamics control the activity and nuclear localization of MRTF/SRF in both compartments, cytoplasm and nucleus, while its target gene functions frequently control cytoskeleton remodelling (15). Interestingly, it has been shown that many MRTF target genes are shared with the activation of YAP/TAZ transcriptional activity (16).

The Yes-associated protein (YAP)/transcriptional coactivator with PDZ-binding motif (TAZ), is a key component of the Hippo pathway and its activity is dependent on Hippo signalling. To stimulate gene expression, YAP/TAZ translocates to the nucleus to interact with transcription factors such as the TEA domain (TEAD) family of transcription factors (17). This process is regulated by kinase signalling in the Hippo pathway. When the Hippo pathway is activated, phosphorylated mammalian STE20-like kinase 1/2 (MST1/2) and Mitogen-activated protein kinase kinase kinase kinases (MAP4Ks) phosphorylate the large tumour suppressor kinase 1/2 (LATS1/2), which promotes YAP/TAZ phosphorylation (18, 19). Subsequently, the 14-3-3 protein binds to phosphorylated YAP/TAZ to retain YAP/TAZ in the cytoplasm, thereby inhibiting YAP/TAZ activity (20). Conversely, when the Hippo pathway is inactive, YAP/TAZ translocates to the nucleus to activate transcription. Hippo signalling is regulated by extracellular matrix stiffness (21), cell density (22), mechanical force (23), and actin dynamics (24). Actin dynamics has been reported to regulate YAP transcriptional activity through the Hippo pathway or the ARID1A–SWI/SNF complex (21, 25, 26).

YAP/TAZ is a master regulator of tissue development and cancer progression, that controls organ size, growth and architecture (20), cell pluripotency, confluency and adhesion (19, 27, 28) but also tumor growth and metastasis (29, 30). Notably, dysregulation of YAP/TAZ has been reported in different tumors (19). YAP is frequently overexpressed in cervical cancer, and expression levels as well as nuclear accumulation are elevated in advanced-stage patients (31). In addition, YAP/TAZ is involved in the different stages of breast cancer, contributing to tumorigenesis, metastasis and therapy resistance (32). Moreover, PanCancer analysis indicates that YAP1 gene fusions involving the N-terminal region of YAP1 are putative oncogenic drivers (33–35). These YAP1-fusion proteins exhibit a constitutive nuclear localization driven by the nuclear localization sequence provided by the fused partner protein, suggesting important functional protein-protein interactions for YAP in the nuclear compartment (36). Although nuclear actin dynamics has been implicated in transcriptional regulation (14), whether actin interacts with YAP in the nuclear compartment for YAP-dependent transcription remained unknown.

Here, we characterize YAP as a G-actin binding protein and identify two G-actin binding regions containing three critical actin binding sites. We show that the subcellular distribution and transcriptional activity of YAP are modulated by the levels of nuclear actin and nuclear G-actin binding to YAP. Moreover, we find out that nuclear G-actin binding to YAP facilitates the formation of YAP–TEAD complexes on DNA. Overall, our results demonstrate that dynamic nuclear YAP-actin interactions regulate YAP signalling for transcription and reveal a novel mechanism in which nuclear actin binding is necessary for YAP–TEAD transcription.

## MATERIAL AND METHODS

### DNA constructs

pET-41 GST-Rpel (MRTF-A) and pRLTK-Renilla-Luc were a gift from Treisman Richard (14). pCMV-flag YAP2 5SA was a gift from Kunliang Guan (Addgene plasmid # 27371; http://n2t.net/addgene:27371; RRID:Addgene_27371) (20). pGL3b-8xGTIIC-luciferase was a gift from Stefano Piccolo (Addgene plasmid # 34615; http://n2t.net/addgene:34615; RRID:Addgene_34615)(21). pEF Flag NLS actin was a gift from Dr. Guido Posern (37). pEGFP-C3-hYAP1 was a gift from Marius Sudol (Addgene plasmid # 17843; http://n2t.net/addgene:17843; RRID:Addgene_17843)(38). pLKO1-shYAP1 was a gift from Kunliang Guan (Addgene plasmid # 27368; http://n2t.net/addgene:27368; RRID:Addgene_27368)(39). EGFP-actin was a gift from B. Imhof (University of Geneva). pmCherry exportin 6 was from the previous work in the lab(40).

pLKO1-shCT was cloned using pLKO1-shYAP as the template, via Gibson assembly with primer o1 and o2 (see Table 1 Primer List). To clone pET28a-TEAD4, TEAD4 coding sequence was amplified from total RNA from HeLa cells using primer o3 and o4, and subclone to pET28a vector via SalI and NotI. pET28a YAP, pET28a YAP NT, pET28a YAP CT and pEF myc YAP was constructed using restriction enzymes. To generate pET28a YAP, pET28a YAP NT, and pET28a YAP CT, YAP sequences were amplified from pEGFP-C3-hYAP1 with primer o5+o6, o7+o8 or o9+o10 then subcloned into pET28a vector via SalI and NotI. To build up pEF myc YAP, YAP sequences were amplified from pEGFP-C3-hYAP1 with primer o29+o30 then subcloned into pEF vector via SpeI and NotI. pET28a YAP N1, pET28a YAP N2, pEF YAP S5A, pWpXL Flag 3NLS EGFP YAP, pEF NLS-actin-YAP was cloned via Gibson assembly. In detail, to clone pET28a YAP N1, pET28a YAP was used as backbone. Two fragments, amplified with primer o11+o14 and o12+o13 were assembled to obtain the indicated constructs. To clone pET28a YAP N2, pET28a YAP NT was used as backbone. Two fragments, amplified with primer o15+o14 and o16+o13 were assembled to obtain the indicated constructs. To build up pEF YAP S5A, fragment containing all 5 single mutations was amplified from pCMV-flag YAP2 5SA with primer o35+o36 and fragment containing the rest of YAP and pEF backbone was amplified from pEF myc YAP with primer o37+o38. Two fragments were applied for Gibson assembly, followed by single mutation mutagenesis with primer o39. To assemble pWpXL Flag 3xNLS EGFP YAP, fragment containing pWpXL flag 3xNLS was amplified from pWpXL flag 3xNLS actin, previously cloned by the lab, with primer o40+o41 and fragment containing EGFP YAP was amplified from pWpXL EGFP-YAP with primer o42+o43, followed by Gibson assembly with both fragments. To construct pEF NLS-actin-YAP, three fragments were amplified and subjected to Gibson assembly. The first fragment containing pEF backbone was amplified from pEF vector with primer o45+o44. The second fragment containing the YAP sequence was amplified from pEF myc YAP with primer o48+o49. The third fragment containing NLS-actin was amplified from pEF NLS-actin with primer o46+o47. To construct single mutants of pET28a YAP, pEF YAP, pEF YAP S5A and pEF NLS-YAP, single mutation mutagenesis was performed with primer o19-o28. To construct pEF YAP DDY, pEF NLS-YAP DDY, pEGFP YAP DDY and pEF NLS-actin-YAP DDY, two rounds of single mutation mutagenesis were performed with primer o26 and o28. To construct YAP shRNA resistant YAP construct, single mutation mutagenesis was performed with primer o50. All generated plasmids were sequence-verified.

### Cell lines

HeLa cell line was a gift from Prof. Norbert Klugbauer and MDA-MB-231 cell line was a gift from Raphael Reuten. HeLa shYAP cells and HeLa shCT cells were generated using lentivirus. pLKO1-shCT or pLKO1-shYAP were transfected together with lentiviral packaging plasmids psPAX2 and pMD2.G into HEK293T cells (ATCC, CRL-3216) to produce virus. HeLa cells were infected with the lentivirus and selected in 1 µg/mL puromycin. HeLa cells and HEK293T cells were maintained in DMEM (Anprotec, AC-LM-0013), w: 4.5 g/L Glucose, w: stable Glutamine, w: Sodium pyruvate, w: 3.7 g/L NaHCO3 supplemented with 10 % fetal constance (FCS, Anprotec, AC-SM-0190), and 100 units/ml penicillin/streptomycin (P/S) (Anprotec, AC-AB-0024). MDA-MB-231 cells were maintained in DMEM/F-12, GlutaMAX™ supplement (Thermo Fisher Scientific, 31331093) supplemented with 10 % fetal constance and 100 units/ml P/S. All cell lines were regularly tested for mycoplasma contamination.

### Protein purification

pET-28a of MRTF-N, TEAD4, and different YAP constructs were transformed into BL21-CodonPlus cell. When the absorbance of 600 nm reached 0.6-0.8, 1 mM IPTG (Roth, CN08.3) was added to induce protein expression and incubated for 16 hours. Pellet was centrifuged at 4, 000 rpm for 20 min and lysed by sonication in TBS buffer (20 mM Tris-HCl, 150 mM NaCl, 1 mM β-mercaptoethanol). After 1 h of 16, 000 rpm centrifugation, the lysate was applied to Ni-IDA Resin (Macherey-Nagel, 745210.120) for purification. Actin protein (>99% pure) from rabbit skeletal muscle (cytoskeleton, Inc.) was purchased from tebubio.

### Size exclusion chromatography

The analytical size-exclusion chromatography was performed on an ÄKTA system (GE Healthcare). G-buffer (5 mM Tris-HCl, pH 7.5, 0.2 mM CaCl_2_, 0.2 mM ATP, 0.5 mM DTT) was used as running buffer. 500 µL of 3 µM YAP, 3 µM G-actin or a mixture of 3 µM YAP and 3 µM actin was injected into Superdex 200 Increase 10/300 GL (Cytiva). Fractions were collected, and mixed with 5x Laemmli sample buffer. After boiling at 95 °C for 10 min, 30 µL of Lysate was loaded onto 10% sodium dodecyl sulfate-polyacrylamide gel electrophoresis (SDS-PAGE) for separation and subjected to immunoblotting.

### *In vitro* F-actin co-sedimentation assay

*In vitro* F-actin co-sedimentation assay was described previously(41, 42). To examine YAP/F-actin binding affinity, 100 µL of 10 µM actin, 5 or 10 µM YAP or a mixture of 10 µM actin and 5 or 10 µM YAP was incubated in 5 mM Tris-HCl pH 7.5, 100 mM KCl, 1 mM MgCl_2_, 0.2 mM CaCl2, 0.2 mM EGTA, 0.2 mM ATP and 0.5 mM DTT for 1 hour at room temperature (RT). After 30 min of ultracentrifugation at 80, 000 x g, 90 μl of supernatant was kept as the non-pelleted fraction (S) and boiled in 22.5 μl of 5× Laemmli buffer. The pellet (P) was washed three times with F-buffer then resuspended and boiled in 125 μl of 1× Laemmli buffer. 20 μl of each sample was loaded into the SDS–PAGE gel and stained with InstantBlue (Abcam, ab119211). Images were acquired with BioRad gel doc EZ Imager equipped with Image Lab software.

### Proximity ligation assay (PLA) and analysis

The proximity ligation assay (PLA) was performed using Duolink in situ Far Red kit reagents (Sigma-Aldrich, DUO92013) according to the manufacturer’s instructions. HeLa wild-type (wt) or HeLa-shYAP cells were seeded onto fibronectin-coated glass coverslips and cultured under sparse conditions. After 24 hours, cells were fixed at room temperature for 10 minutes with 4 % paraformaldehyde (PFA) in phosphate-buffered saline (PBS), followed by permeabilization in 0.3 % Triton X-100 in PBS. To block non-specific antibody binding, samples were incubated in 3 % bovine serum albumin (BSA) in PBS for 30 minutes at 37 °C in a pre-heated humidity chamber. Primary antibodies were diluted in a buffer containing 3 % BSA and 0.05 % Tween-20 in PBS. For co-detection experiments, both antibodies were combined in the same buffer. Following the removal of the blocking solution, the primary antibody mix was applied to each sample and incubated overnight at 4 °C. After gentle PBS washes, PLA probes (anti-mouse MINUS, DUO92004 and anti-rabbit PLUS, DUO92002, Sigma-Aldrich) were diluted in the same antibody buffer and added to the samples. Probe hybridization was carried out for 1 hour at 37 °C in a humidity chamber. Post-hybridization, slides were washed twice with 1× Wash Buffer A (0.01 M Tris, 0.15 M NaCl, 0.05% Tween-20; pH 7.4) for 5 minutes under gentle agitation. The ligation solution was prepared by diluting ligation stock 1:5 in high-purity water, and ligase was freshly added at a final dilution of 1:40 immediately prior to sample application. Samples were incubated with the ligation mixture for 30 minutes at 37 °C. Rolling circle amplification was performed by diluting amplification stock 1:5 in high-purity water, with polymerase added at a final dilution of 1:80. Amplification was carried out in the dark for 100 minutes at 37 °C in a pre-heated humidity chamber due to the light sensitivity of the reagents. The following primary antibodies were used: rabbit polyclonal anti–β-actin (1:50; Abcam, ab8227), mouse monoclonal anti–YAP (1:50; Abnova, H00010413-M01), and rabbit monoclonal anti–Tead1 (1:50; Cell Signaling Technology, 12292S).

Images were acquired using a Zeiss LSM800 confocal laser-scanning microscope equipped with a 63×/1.4 NA oil objective, DAPI and Cy5 filter sets, Zen Blue software, and Airyscan Multiplex SR-4Y super-resolution mode. For each sample, Z-stacks comprising 35 optical sections (0.14 μm step size) were collected with DAPI and Cy5 channels. Image analysis was conducted using IMARIS v10.1.0 (Oxford Instruments). Nuclei were rendered in 3D based on the DAPI channel and subsequently masked using the surface masking tool. Cy5-labeled PLA signals were visualized and quantitatively assessed within nuclear and cytoplasmic compartments across all experimental conditions.

### Alphafold prediction

The prediction of the interaction of G-actin (UniProt P68133) with YAP (UniProt P46937) was performed with AlphaFold 3 (43). The prediction of the interaction of G-actin (UniProt P68133) with YAP-N was performed with Alphafold2-multimer(44–46). The binding interface was identified upon the presence of hydrogen bonds shorter than 3.5 Å. Plddt plot was generated using af_plotter.py (https://github.com/LMSBioinformatics/af_plotter). The hydrogen bond identification and the structure editing were conducted using Pymol.

The prediction of the structure of TEAD4 (UniProt Q15561)/M-CAT DNA, YAP/TEAD4/M-CAT DNA and YAP/G-actin/TEAD4/M-CAT DNA was performed with AlphaFold 3(43). The structure editing was conducted using Pymol (Schrodinger, LLC). The movie was generated using Openshot video editor.

### ChIP procedure

Chromatin-Immunoprecipitations (ChIP) were performed using the Pierce Magnetic ChIP Kit (Thermo Fisher Scientific, 26157) according to the manufacturer’s protocol with optimization for crosslinking(47, 48). In brief, cells were crosslinked with 2 mM disuccinimidyl glutarate (Thermo Fisher Scientific, 20593) for 20 minutes, followed by 1 % formaldehyde (Thermo Fisher Scientific, 410731000) for an additional 10 minutes. The crosslinking was stopped by adding glycine (component in Pierce Magnetic ChIP Kit). After removing the medium, cells were washed twice with ice-cold PBS and collected with scraping in ice-cold PBS containing Halt Cocktail (component in Pierce Magnetic ChIP Kit) followed by 5-minute centrifugation at 3000 x g. Cell pellets were lysed with membrane extraction buffer (component in Pierce Magnetic ChIP Kit) and nuclei were collected by centrifuging for 3 minutes at 9000 x g.

Nuclei were resuspended in MNase Digestion Buffer (component in Pierce Magnetic ChIP Kit) and digested with MNase (component in Pierce Magnetic ChIP Kit) for 15 minutes at 37 °C. Digestion was stopped by adding MNase Stop Solution (component in Pierce Magnetic ChIP Kit) and incubating on ice for 5 minutes. The nuclei were recovered by 5-minute centrifugation at 9000 x g, resuspended in IP dilution buffer (component in Pierce Magnetic ChIP Kit) and subjected to sonication with a Bioruptor (Diagenode) at high amplitude for 8 × 30 s:30 s [on:off cycles]. After centrifuging at 9000 x g for 5 minutes, DNA concentration in the chromatin mixture was measured by Qubit Fluorometer (Thermo Fisher Scientific) with Qubit™ dsDNA Quantification Assay Kits (Thermo Fisher Scientific, Q32854). Chromatin containing 6 µg dsDNA was kept as 10 % input and chromatin containing 60 µg dsDNA was incubated with the indicated antibody at 4 °C for 2 hours. Added CHIP grade protein A/G magnetic beads (component in Pierce Magnetic ChIP Kit) to the supernatant and incubated it at 4 °C for 2 hours. Collected the beads with a magnetic stand, removed the supernatant and washed the beads three times with IP wash buffer 1 (component in Pierce Magnetic ChIP Kit) and one time with IP wash buffer 2 (component in Pierce Magnetic ChIP Kit). Immunoprecipitated chromatin on the magnetic beads was eluted in IP elution buffer (component in Pierce Magnetic ChIP Kit) with 30-minute incubation at 65 °C, followed by Proteinase K (component in Pierce Magnetic ChIP Kit) digestion with 90-minute incubation at 65 °C. DNA was recovered with the DNA clean-up kit (component in Pierce Magnetic ChIP Kit).

### RNA isolation, reverse transcription and quantitative PCR

HeLa cells were seeded in a 6-well plate were transfected after 24 hours with the indicated constructs using FuGENE® HD Transfection Reagent (Promega, E2311). After 48-hour incubation, cells were lysed with Lysis Buffer RLT (component in RNeasy Kits for RNA Purification) directly in the 6-well plate and total RNA was extracted from each sample using the RNeasy Kits for RNA Purification (Qiagen, 74106). The concentration of purified RNA was measured with a NanoDrop spectrophotometer (Thermo Fisher Scientific). Reverse transcription was performed in two steps, sample denaturation and complementary DNA (cDNA) synthesis. RNA and primer were denatured in a 10 µL reaction with 1 µg total RNA, 2 µL Oligo d(T)23 VN (New England Biolabs, S1327S), 1 µL 10 mM dNTP (New England Biolabs, N0447S) for 5 min at 65 °C, then cDNA synthesis was synthesized at 42 °C for 1 hour after adding 2.8 µL Nuclease-free H_2_O (New England Biolabs, B1500L), 4 µL 5X ProtoScript II Buffer, 2 µL 0.1M DTT (Roth, 6908.1), 0.2 µL RNase Inhibitor (40 U/μl) (New England Biolabs, M0307S) and 1 µL ProtoScript® II Reverse Transcriptase (New England Biolabs, M0368L). Quantitative PCR (qPCR) for gene expression and CHIP-qPCR analysis was performed in a 20-µl reaction mixture containing 2 µl cDNA or recovered DNA from CHIP, 200 nM primers, and 10 µl GoTaq® qPCR Master Mix (2X) (Promega, A6002).

For relative gene expression analysis, the expression levels of target genes were normalized to those of the internal control gene, GAPDH, by calculating the ratio of their expression levels (R), using equation R=2 ^ (Cq_GAPDH_-Cq_target gene_) x 100. In the ChIP-qPCR experiments, bound DNA levels (BD) were assessed by comparing ChIP samples to a 10 % input using equation BD = 2 ^ (Cq_10%input_-Cq_sample_-3.32) x 100. Each reaction was performed in technical triplicates, and three biological replicates were used for each condition. Primers used for qPCR are from previous research (16, 49) and listed in Supplementary Table S2.

### SPR – Surface Plasmon Resonance measurements

Dynamic interaction of > 99% rabbit muscle actin (Cytoskeleton) or TEAD4 with protein ligands were analyzed by using Biacore X100 system (GE Healthcare). Two lanes on a CM5-chip were generated by coupling 10 μg/ml actin or TEAD4 covalently to one lane using 400 mM N-ethyl-N-dimethylaminpropyl-carbodiimide (EDC) and 100 mM N-hydroxy-succinimide (NHS). Afterwards both lanes were saturated with 1 M ethanolamide resulting in an actin-coated or TEAD4-coated (lane 2) and a control lane (lane 1) for exclusion of non-specific binding. In general ligand partners were guided over both lanes. G-actin or TEAD4 was immobilized on the surface of a CM5 chip (Cytiva, 29149604) according to the built-in program. Briefly, G-actin was dialyzed in a buffer containing 5 mM HEPES (Roth, 3105.3) pH 7.4, 0.2 mM CaCl_2_ and 0.2 mM ATP and TEAD4 was dialyzed in 10 mM HEPES pH 8.8, 150 mM NaCl. To immobilize G-actin or TEAD4, one lane on a CM5 chip was activated with a 1:1 mixture of 400 mM N-ethyl-N-dimethylaminpropyl-carbodiimide (EDC) and 100 mM N-hydroxy-succinimide (NHS). G-actin or TEAD4 was diluted in 10 mM sodium acetate buffer pH 4.5 to a final concentration of 1 µM and then injected onto the chip to reach a target level of 1000 resonance units (RU). Remaining activated carboxyl groups were quenched by injecting 1 M ethylenediamine-HCl at pH 8.5. For each recombinant protein, five different concentrations of recombinant proteins and a buffer-only control were injected sequentially for 180 s and followed by a 600 s dissociation. The running buffer for actin binding analysis contained 10 mM HEPES pH 7.4, 150 mM NaCl, 3mM EDTA, 10 mM MgCl_2_, 0.2 mM ADP, 0.5 mM DTT and 0.05% Tween-20 (Roth, 9127.1) and for TEAD4 binding assay contained 10 mM HEPES pH 8.8, 150 mM NaCl, 3mM EDTA and 0.015% Tween-20. Regeneration between measurements was achieved using 10 mM NaOH. Each measurement was repeated at least three times. All binding sensorgrams were collected, processed using the Biacore X-100 Evaluation software and analyzed with the Prism 10 (GraphPad Software, LLC. (Version 10.5.0). Bound protein was determined as relative response units (RU) corrected for the unspecific binding to lane 1 (RULane2-RULane1). Kinetic constants (K_D_) were calculated by plotting of RU of steady state versus analyte concentration.

### Immunoblotting and antibodies

To verify successful YAP silencing or the transfection of indicated constructs, cells were transfected with the respective plasmid and lysed after 48 hours by adding 200 µL/well of 2x Laemmli sample buffer directly into a 6-well plate. After boiling at 95 °C for 10 minutes, 30 µL of Lysate was loaded to 10 % SDS-PAGE gel for separation. Proteins were then transferred to a PVDF membrane (Cytiva, 10600022) using the Power Blotter XL System (Invitrogen) with a semi-dry transfer method. The PVDF membranes then were blocked with 5% milk for 1 hour at RT, incubated with primary antibodies overnight at 4 °C and followed with 1-hour incubation with a Horseradish peroxidase-labeled secondary antibody at RT. Proteins were detected using Amersham Imager 600 (GE Healthcare Life Sciences) with SuperSignal West Femto Maximum Sensitivity Substrate (Thermo Fisher Scientific, 34096). The following primary were used: mouse monoclonal anti-FLAG® M2 antibody (1:1000, Merck, F1804), YAP (D8H1X) XP® Rabbit mAb (1:500, Cell Signaling Technology, 14074), rabbit polyclonal anti-Myc Tag (1:1000, Cell Signaling Technology, 2272), mCherry (E5D8F) Rabbit mAb (1:1000, Cell Signaling Technology, 43590), anti-Tubulin (11H10) Rabbit mAb (1:1000, Cell Signaling Technology, 2125), mouse monoclonal anti-Actin antibody (Merck, A4700).

### Fluorescence recovery after photobleaching (FRAP)

HT1080 cells were seeded on 2cm glass bottom dishes and transfected with GFP-actin (). 24 hours after transfection, FRAP experiments were performed in LSM 800 confocal laser scanning microscope (Zeiss) equipped with a 63×/1.4 NA oil objective. For GFP-actin bleaching, a 4 µm diameter circular region of interest (ROI) was bleached with 15 iterations of 100% laser power at 488 nm. Images were acquired every second for a total of 60 cycles and collected at a 12-bit intensity resolution of 256×256 pixels with a pixel dwell time of 2.06 µs. Photobleaching initiated after three images were taken. Fluorescence intensities from nuclear and cytosolic ROIs were collected from Zen blue (3.4), normalized to the maximum intensity for every ROI and plotted in GraphPad (10.5.0).

### Immunofluorescence

HeLa cells were seeded in sparse or confluent conditions on the fibronectin (Merck, F1141)-coated coverslips and transfected with indicated plasmids. After 48 hours, cells were washed with 1 X PBS, fixed with 4 % paraformaldehyde solution in PBS (SantaCruz, sc-281692) for 10 min and permeabilized using 0.3 % Triton X-100 (Roth, 3051.2). After 1 hour blocking in 5 % FCS, coverslips were incubated with primary antibody overnight at 4 °C, followed by 1 hour incubation of secondary antibody at RT. The coverslips were mounted onto glass slides using ProLong Diamond Antifade Mountant (Thermo Fisher Scientific, P3970). To observe distribution of the endogenous YAP with the overexpression of NLS-actin or mCherry-exportin 6, images were acquired using a Zeiss LSM800 confocal laser-scanning microscope in confocal mode, employing a 63×/1.4 NA oil objective with a z-stack interval of 0.22 µm.

For quantifying the nuclear-to-cytoplasmic (nuc/cyto) ratio of YAP^WT^ or YAP^DDY^, images were obtained using the same settings, with a z-stack interval of 0.5 µm. Image analysis was conducted using IMARIS v10.1.0 (Oxford Instruments). To analyze the YAP nuc/cyto ratio (Fig. 4D-F), nuclei were rendered in 3D based on the DAPI channel and subsequently masked using the surface masking tool. Intensities of YAP within nuclear and cytoplasmic compartments were quantified and used to calculate the ratio.

The following primary and secondary antibodies were used: mouse monoclonal anti-FLAG® M2 antibody (1:500, Merck, F1804), YAP (D8H1X) XP® Rabbit mAb (1:50, Cell Signaling Technology, 14074), rabbit polyclonal anti-Myc Tag (1:200, Cell Signaling Technology, 2272), chicken anti-Rabbit IgG (H+L) Cross-Adsorbed Secondary Antibody, Alexa Fluor™ 488 (1:400, Thermo Fisher Scientific, A-21441), Chicken anti-Mouse IgG (H+L) Cross-Adsorbed Secondary Antibody, Alexa Fluor™ 488 (1:400, Thermo Fisher Scientific, A-21200), goat anti-Rabbit IgG (H+L) Cross-Adsorbed Secondary Antibody, Alexa Fluor™ 647 (1:200, Thermo Fisher Scientific, A-21244), and 4′, 6-diamidino-2-phenylindole (DAPI).

### 8X GTIIC-luciferase gene reporter assay

HeLa cells or MDA-MB-231 cells were seeded into a 6-well plate. After 24 hours, cells were transfected with 200 ng of pRLTK-Renilla-Luc, 400 ng of pGL3b-8xGTIIC-luciferase and 500 ng of the indicated plasmids per well. FuGENE® HD Transfection Reagent (Promega, E2311) was used for HeLa cells and Lipofectamine 3000 (Thermo Fisher Scientific, L3000015) was used for MDA-MB-231 cells as transfection reagent. After a 48-hour incubation, the cells were lysed with 200 µL Triton lysis buffer (0.15 M Tris, 75 mM NaCl, 3 mM MgCl_2_, 0.25% Triton X-100) for 10 minutes on a shaker at 4 °C. Cells were then scraped and centrifuged at 16, 000 x g and 4 °C for 20 minutes. For luminescence measurements, 100 µl of each lysate was pipetted into a 96-well plate and 50 µl of firefly buffer (15 mM DTT, 0.6 mM coenzyme A, 0.45 mM ATP, 4.2 mg/ml d-luciferin) was added simultaneously to the lysates. Luminescence was measured immediately. Subsequently, 75 µl of renilla buffer (45 mM EDTA, 30 mM Na_4_P_2_O_7_, 1.425 M NaCl, 0.06 mM PTC124, 0.01 h-CTZ) was added to the reaction and luminescence was measured again. The ratio of luminescence between luciferase and renilla was calculated and used as the relative luciferase activity, or normalized against the average of the indicated control. Moreover, 50 µL of each lysate was mixed with 12.5 µL 5x Laemmli sample buffer, boiled at 95 °C for 10 minutes and subjected for immunoblotting (Extended Data Fig. 3A-G).

### Cell proliferation assay

The proliferation assay for HeLa cells was conducted using the Click-iT™ Plus EdU Alexa Fluor 647 Imaging Kit (Thermo Fisher Scientific, C10640). HeLa cells were seeded onto a fibronectin-coated coverslip in a 6-well plate. After 24 hours, cells were transfected with 500 ng of pEF, pEF-YAP^WT^ or pEF-YAP^DDY^ using FuGENE® HD Transfection Reagent. Following a 48-hour incubation, cells were treated with 5 µM EdU for 4 hours, then fixed and subjected to immunostaining. Images were captured using a Zeiss LSM800 confocal laser-scanning microscope, utilizing a 40×/1.3 NA oil objective. The proliferation ability was indicated by calculating the ratio of EdU-positive transfected cells to the total number of transfected cells.

### Statistical analysis

Statistical analyses and graphical representations were carried out using GraphPad/Prism v.10.5.0. Information on exact n values, statistical tests and their description, and error bars are given in the figure legends. In general, experimental data sets were tested for normality. Statistical significance was evaluated with one-way ANOVA for multiple comparisons (Tukey’s). For ChIP-qPCR and mRNA expression experiment, a two-tailed t-test was used. Data are presented as either column bar graph ± s.d., scatter dot bar plot with mean ± s.d. or violin plots showing all data points with median and quartiles. Statistical significance is indicated as *P < 0.05, **P < 0.01, ***P < 0.001, ****P < 0.0001.

## RESULTS

### YAP binds G-actin

To examine a potential relationship between YAP and nuclear actin, we initially performed proximity ligation assays (PLA), which allow for endogenous detection of protein interactions within vicinities of less than 40 nm (50). We had previously optimized PLA detection for endogenous nuclear proteins (51). Using this assay, we found that YAP and actin robustly interacted comparable to YAP and TEAD1 (Fig. 1a, b, c), and that this interaction was dependent on actin, since silencing of YAP using shRNA abolished detectable PLA dots (Fig. 1a, b, c). To further characterize the subcellular localization of YAP–actin interactions, we quantified the PLA signal distribution across cellular compartments. Analysis revealed that 52% of PLA dots were located in the cytoplasm, while 48% were detected in the nucleus (Extended Data Fig. 1a), indicating that the interaction between YAP and actin occurs in both compartments.

**Figure 1.**
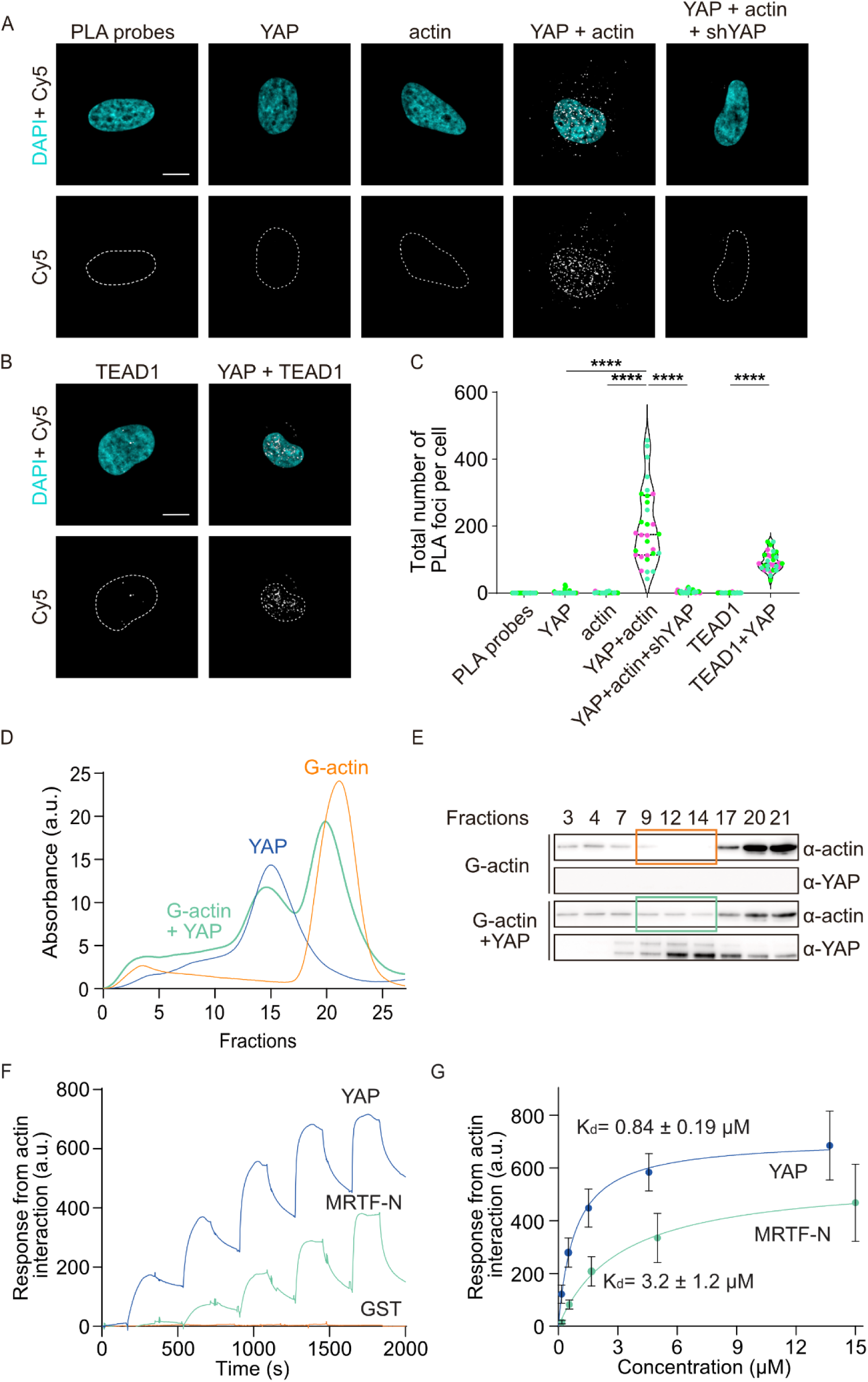
The transcription factor YAP interacts directly with actin. (A-B) Representative images of HeLa cells subjected to Proximity Ligation Assay (PLA) with the indicated primary antibodies. Nucleus is indicated by white dashed line. Scale bar = 10 μm. Images are shown as maximum intensity projection (MIP). (C) Violin plot with individual data points (green, magenta, cyan), median and quartiles showing quantification of total number of PLA puncta. Data is n = 30 cells over three independent biological replicates (D) Size-exclusion chromatography of conditions in the presence of YAP only (blue), G-actin only (orange) and G-actin+YAP (green). (E) Immunoblot analysis of indicated conditions and fractions from (D), orange box indicates fractions excluded for G actin, green box indicates YAP/actin complex-containing fractions. (F) Representative sensorgrams of surface plasmon resonance (SPR) obtained from injections of purified YAP (blue), MRTF-N (green) and GST (orange) over G-actin-coated CM5 chip. (G) Dose-response curve of SPR for purified YAP (blue) and MRTF-N (green) indicating the dissociation constant (K_d_). Data are mean ± SEM over three independent biological replicates. Statistical analysis was done using One-way ANOVA with Tukeýs multiple comparison test ****p < 0.0001.

To characterize the dynamics of the nuclear actin pool involved in YAP interaction, we overexpressed GFP-actin in HT1080 cells and performed FRAP experiments in the cytoplasm or nucleus. Both nuclear and cytosolic GFP-actin showed comparable and rapid recovery kinetics (Extended Data Fig. 1b, c), indicating that actin in the nucleus is maintained as a freely available highly mobile pool that exhibits similar dynamics as actin in the cytosol.

Given that actin dynamics regulates MRTF-A activity through dynamic G-actin/RPEL interactions (13), we investigated whether YAP might exhibit similar properties. For this, we combined protein reconstitution assays with size-exclusion chromatography (SEC), as well as F-actin co-sedimentation assays in order to investigate G- and or F-actin interactions. Compared to the SEC curve of actin or YAP alone, the SEC curve in the presence of YAP and actin showed an increased absorbance of fractions 9-14 and a decreased absorbance of YAP corresponding fractions (14–17), with actin corresponding fractions (20–23) (Fig. 1d). Western blot analysis of the SEC fractions showed that actin was only detected from fractions 9-14 in the presence of YAP and actin but not in the presence of actin alone indicating YAP binding to G-actin (Fig. 1e), whereas the modest decrease of the absorbance of actin and YAP eluents on the SEC curve suggests that the interaction might be transient or of a more dynamic nature. In contrast, the presence of F-actin does not co-sediment with increasing amount of YAP (Extended Data Fig. 1d), indicating that YAP does not interact with F-actin.

To determine the binding affinities between YAP and actin or between the MRTF N-terminus (MRTF-N) and actin, we applied surface plasmon resonance (SPR) measurements. MRTF-N, which binds actin (14, 52), was used as a positive control, whereas GST alone served as the negative control. MRTF-N bound actin with a *K*_d_ of 3.2 ± 1.2 μM, in line with previous study (52), while GST does not bind with actin. YAP bound actin with a *K*_d_ of 0.84 ± 0.19 μM, exhibiting higher actin-binding affinity compared to MRTF-N (Fig. 1f, g). Notably, the YAP-actin complex displayed a fast dissociation rate constant k_off_ = 3.781 ± 0.3360 x 10^-3^ s^-1^ and a half-life dissociation t_1/2_ = 184.4 ± 16.21 s (Fig. 1f) indicative of dynamic interactions between YAP and G-actin.

### The YAP N-terminus contains two G-actin-binding interfaces

YAP possesses a TEA domain transcription factor (TEAD) binding domain (TB) at the N-terminus and a C-terminal transcriptional activation domain (TAD) linked by a WW domain (53) (Fig. 2a). Based on these functional domains, we generated two YAP truncation variants YAP-N and YAP-C to measure their binding affinities to actin (Fig. 2a). SPR measurements revealed that YAP-N displayed actin binding affinities comparable to YAP (Fig. 2c), while YAP-C displayed a weaker actin binding affinity (Fig. 2b). To more precisely determine the location within YAP for actin binding, YAP-N was split further into YAP-N1 and YAP-N2 (Fig. 2a), for SPR analysis. Surprisingly, both YAP fragments exhibited actin binding affinity (Fig. 2b, d), indicating that YAP binds to G-actin via two regions within its N-terminal domain.

**Figure 2.**
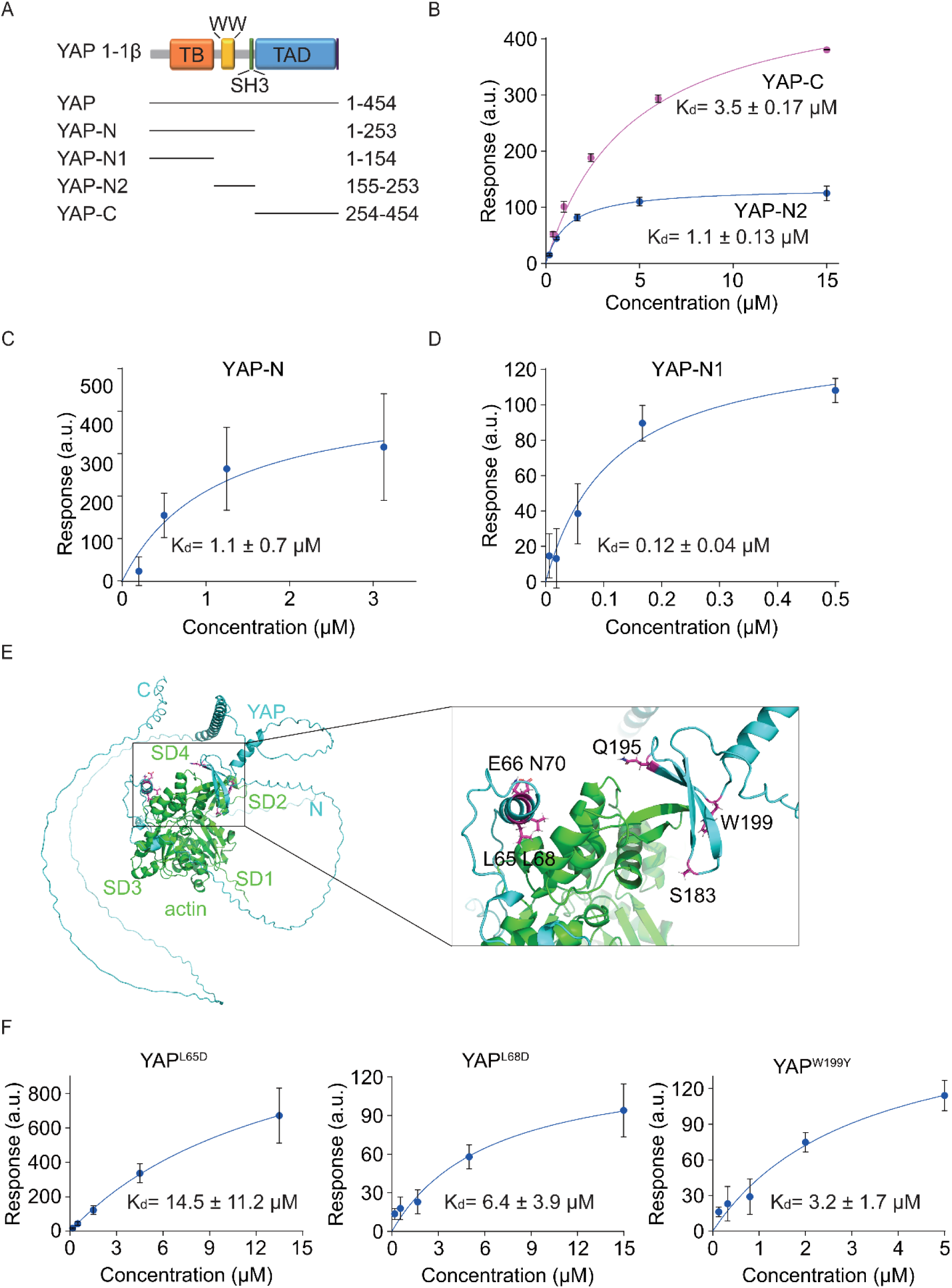
Identification of two actin binding domains located on the N-terminus of YAP. (A) Schematic representation of YAP full length with indicated functional domains, TEAD-binding domain (orange), WW-domain (yellow), SH3-binding domain (green), transactivation domain (TAD, blue), PDZ-binding motif (magenta) and purified YAP truncations. (B–D) Dose-response curves for purified (B) YAP-N2 (blue) and YAP-C (magenta), (C) YAP-N and (D) YAP-N1 with indicated dissociation constant (K_d_). Data are mean ± SEM. Data are from three independent biological experiments, except (B): YAP-C: n = 2. (E) AlphaFold3 prediction model of YAP (cyan) interaction with G-actin (green). Overview (left) and zoom view (black box) of predicted binding interface (right). Predicted binding sites on YAP are shown in magenta and identified amino acids are indicated. (F) Dose-response curves for purified YAP^L65D^ (left panel), YAP^L68D^ (middle panel), and YAP^W199Y^ (right panel) indicating the dissociation constant (K_d_). Data are mean ± SEM. Data are from three independent biological experiments.

### YAP-L65, -L68 and -W199 are involved in YAP actin interactions

To identify potential key YAP residues involved in actin binding, we combined AlphaFold predictions with targeted single-site mutagenesis. YAP and YAP-N were used to simulate actin binding using AlphaFold 3 (43) (Fig. 2e) or AlphaFold 2 (44–46) (Extended Data Fig. 2a, b). Consistent with the results from the YAP truncations (Fig. 2a, b, c, d), two actin binding regions were predicted in the YAP N-terminus, while the C-terminus was not predicted to be implicated in actin binding (Fig. 2e). Seven amino acids (L65, E66, L68, N70, S183, Q195, and W199) were predicted to be involved in actin binding and were subsequently substituted with amino acids exhibiting opposite physicochemical properties but similar molecular weights. Specifically, YAP^L65D^, YAP^E66R^, YAP^L68D^, YAP^N70L^, YAP^S183A^, YAP^Q195L^, YAP^W199Y^ were generated and applied for further SPR analysis (Fig. 2f and Extended Data Fig. 2c). As a negative control, we generated a YAP^Q290L^ mutant (Fig. 2e). Interestingly, YAP^L65D^, YAP^L68D^ as well as of YAP^W199Y^ impaired actin binding (Fig. 2f) consistent with AlphaFold predictions for both, YAP/actin and YAP-N/actin (Fig. 2e and Extended Data Fig. 2a). In contrast, all other tested mutations had comparable actin binding affinities to YAP^WT^ (Extended Data Fig. 2 c, d).

### Nuclear actin modulates YAP subcellular localization and transcriptional activity

Recent studies have highlighted the importance of nuclear actin in transcriptional regulation (11, 42, 54). We therefore set out to determine whether nuclear G-actin binding to YAP is important for its transcriptional activity.

First, we performed immunofluorescence (IF) staining in confluent and sparse cells to determine whether nuclear actin can alter YAP subcellular localization, a process known to correlate with actin dynamics and cell confluence (21, 22, 55). Consistent with previous results (22), YAP was predominantly localized in the nuclear compartment under sparse conditions, while it was primarily distributed in the cytoplasm under confluent conditions (Fig. 3a, b, c). Interestingly, overexpression of a nuclear-localized actin (NLS-actin) in confluent cells relocated YAP to the nucleus (Fig. 3a, c) while depletion of nuclear actin through overexpression of exportin 6 in sparse cells (56) resulted in YAP cytoplasmic retention (Fig. 3a, b), indicating that nuclear actin levels modulate the cellular distribution of YAP.

**Figure 3.**
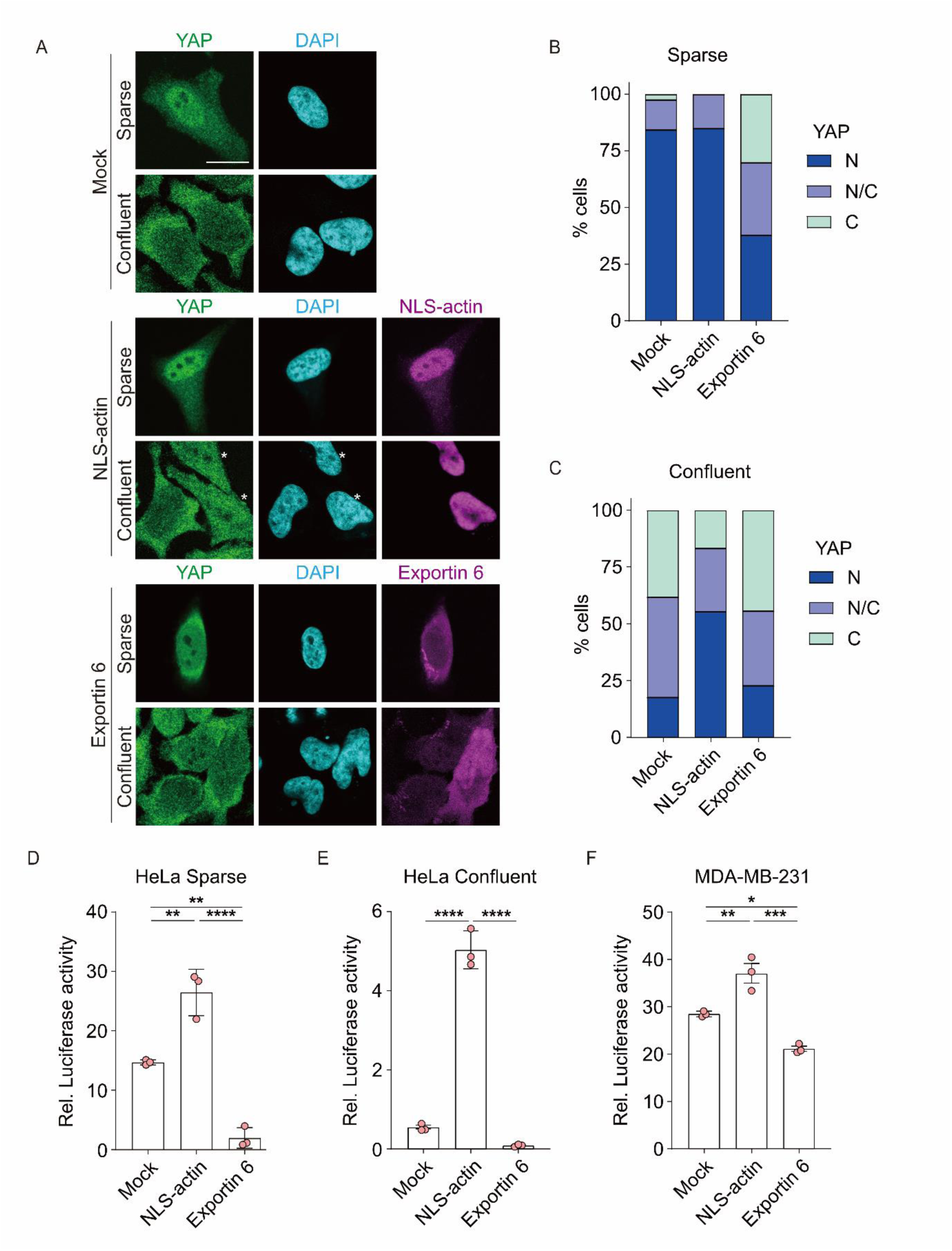
Nuclear actin regulates the subcellular localization and transcriptional activity of YAP. (A) Immunofluorescence of endogenous YAP (green) and DAPI (blue) in HeLa cells in sparse and confluent conditions, transfected with empty vector (Mock, top panel), NLS-actin (middle panel, magenta) or mCherry-exportin 6 (bottom panel, magenta). White asterisks show the NLS-actin-positive cells. Images are shown as full z-stacks with 200 nm z-distance. (B-C) Quantitation of YAP subcellular localization in (B) sparse and (C) confluent conditions (N, blue: nuclear localization, N/C, purple: nuclear and cytoplasmic localization, C, green: cytoplasmic localization). (D-F) Luciferase assay for YAP activity in (D-E) HeLa cells for sparse and confluent conditions or (F) MDA-MB-231 cells, transfected with luciferase plasmids and co-transfected with empty vector (Mock), NLS-actin or mCherry-exportin 6 and quantified 48 h after transfection. Scale bar = 20 μm. Scatter dot bar plot showing data with mean ± SEM from three independent biological experiments. Each dot represents an independent experiment. Statistical analysis was done using One-way ANOVA with Tukeýs multiple comparison test *p < 0.05, **p < 0.01, ***p < 0.001, ****p < 0.0001.

Next, we analysed whether nuclear actin mediates YAP transcriptional activity. For this, we performed 8×GTIIC-lux luciferase reporter gene assays as previously described (21). Using these assays, we could confirm lower YAP activity under confluent conditions versus higher Yap activity in sparse conditions (Fig. 3d, e). Of note, overexpression of NLS-actin significantly enhanced YAP activity, while exportin 6 overexpression robustly reduced YAP activity under both sparse and confluent conditions in HeLa cells, as well as under confluent condition in MDA-MB-231 cells (Fig. 3d, e, f). These results show that nuclear actin levels modulate YAP’s subcellular distribution and transcriptional activity.

### Nuclear G-actin-YAP binding modulates YAP subcellular localization

To determine whether G-actin binding affects the subcellular distribution of YAP, we performed immunofluorescence on cells expressing YAP^WT^ (wild-type YAP) and YAP^DDY^ (YAP^L65D^ ^L68D^ ^W199Y^, a YAP construct harbouring three actin-binding-deficient mutations). Interestingly, in both sparse (Fig. 4a, d) and sub-confluent cells (Fig. 4b, e), YAP^DDY^ displayed a more prominent cytoplasmic distribution when compared to YAP^WT^. However, in confluent cells, YAP^WT^ as well as YAP^DDY^ displayed prominent cytoplasmic distributions (Fig. 4c, f). These results suggest that cell density control of YAP subcellular localization relies on actin/YAP interactions.

**Figure 4.**
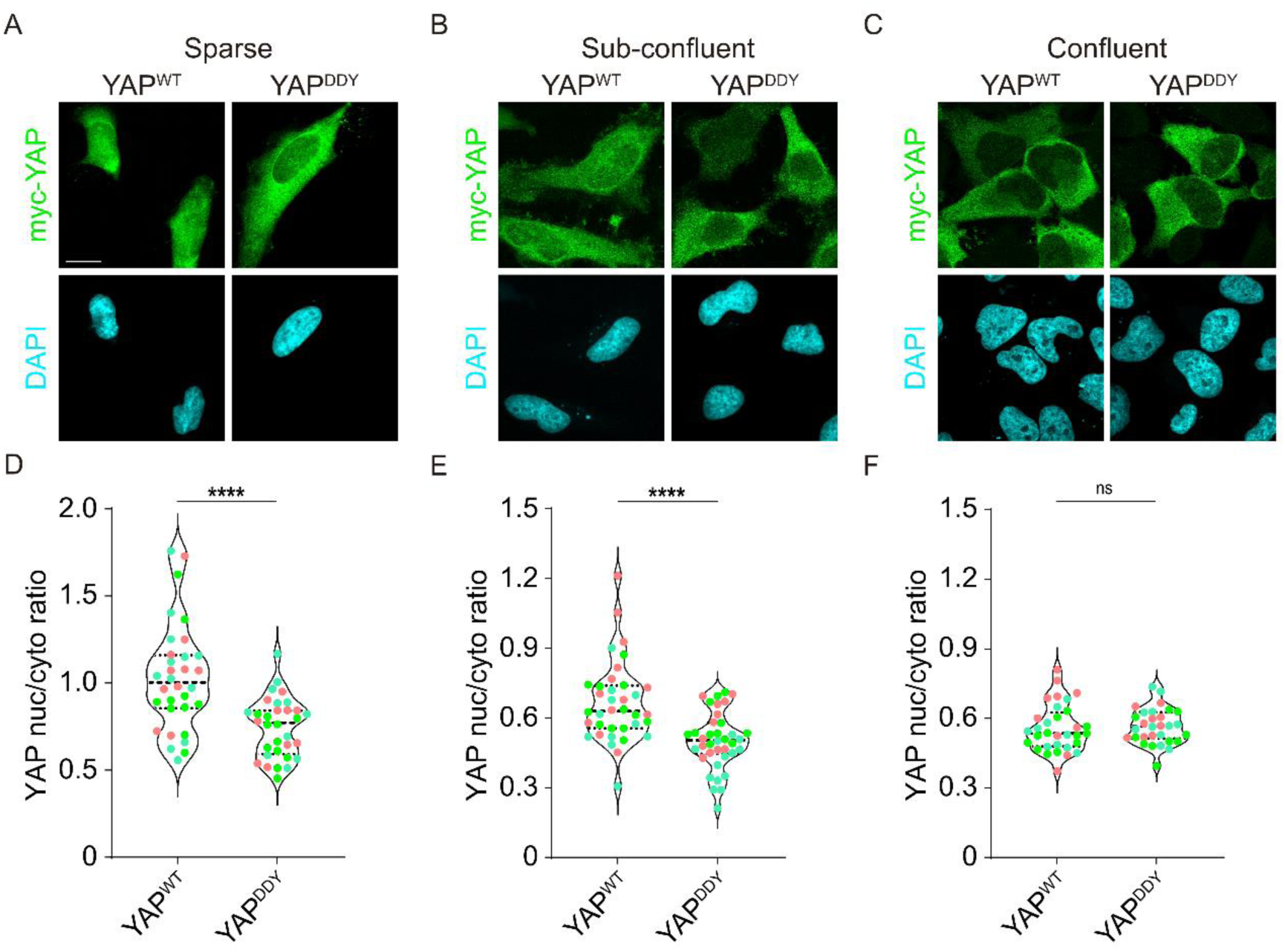
Actin-YAP binding regulates the subcellular localization of YAP. (A-C) Immunofluorescence of myc-YAP (green, top) and DAPI (blue, bottom) in HeLa cells in sparse, sub-confluent and confluent conditions. HeLa cells were transfected with myc-YAP^WT^ or myc-YAP^DDY^. Images are shown as full z-stacks with 500 nm z-distance. Scale bar = 20 μm. (D-F) Violin plot with individual data points (green, pink, cyan), median and quartiles showing the quantification of the nuc/cyto ratio of YAP subcellular distribution. Data are from three independent experiments, n = 32, 32, 36, 38, 32, 30. Each dot represents one cell. Two-tailed unpaired Student’s t-test was used to calculate p-values. *p < 0.05, **p < 0.01, ***p < 0.001, ****p < 0.0001.

### Nuclear G-actin-YAP binding is essential for YAP transcriptional activity

To determine the importance of G-actin binding for YAP function, we compared the transcriptional activities of YAP^WT^ with YAP^DDY^. As expected, expression of YAP^WT^ led to an increased transcriptional activity, whereas YAP^DDY^ exhibited a 50% reduction in transcriptional activity compared to YAP^WT^ (Fig. 5a). To further corroborate this, we performed rescue experiments in a HeLa cell line depleted for YAP using shRNA and assessed transcriptional activity using reporter gene assays. Silencing of YAP effectively reduced YAP activity (Fig. 5b, Extended Data Fig. 3d), and EGFP-YAP^WT^ rescued YAP activity whereas EGFP-YAP^DDY^, deficient in G-actin-binding, did not (Fig. 5b).

**Figure 5.**
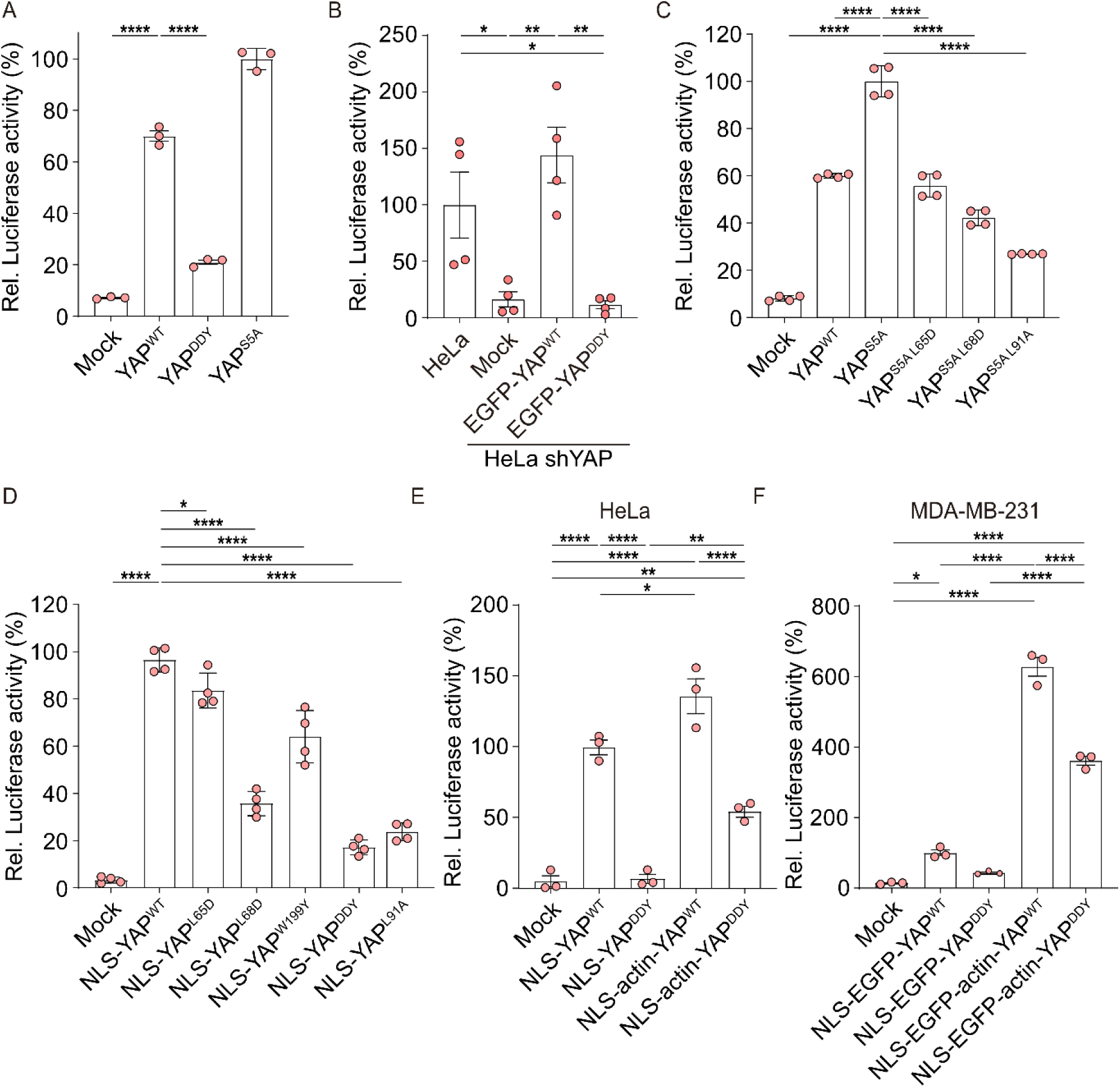
Binding of nuclear actin with YAP mediates its transcriptional activity via regulation of the YAP-TEAD-DNA interaction. (A) Luciferase assay for YAP activity in HeLa cells transfected with luciferase plasmids, co-transfected with empty vector (Mock), YAP^WT^, YAP^DDY^ or YAP^S5A^. The data were normalized to the average luciferase activity of YAP^S5A^. (B) Luciferase assay for YAP activity in HeLa cells and HeLa shYAP cells transfected with luciferase plasmids, co-transfected with empty vector (Mock), EGFP-YAP^WT^ or EGFP-YAP^DDY^. The data were normalized to the average luciferase activity of wt HeLa cells. (C) Luciferase assay for YAP activity in HeLa cells transfected with luciferase plasmids, co-transfected with empty vector (Mock), YAP^WT^ and indicated YAP^S5A^ mutants. The data were normalized to the average luciferase activity of YAP^S5A^. (D) Luciferase assay for YAP activity in HeLa cells transfected with luciferase plasmids, co-transfected with empty vector (Mock), indicated NLS-YAP mutants. The data were normalized to the average luciferase activity of NLS-YAP^WT^ (E-F). Luciferase assay for YAP activity in (E) HeLa cells and (F) MDA-MB-231 cells transfected with luciferase plasmids, co-transfected with empty vector (Mock), indicated (E) NLS-YAP or (F) NLS-EGFP-YAP constructs. The data were normalized to the average luciferase activity of (E) NLS-YAP^WT^ or (F) NLS-EGFP-YAP^WT^. Luciferase activity was quantified 48 h after transfection, statistical analysis was done using One-way ANOVA with Tukeýs multiple comparison test *p < 0.05, **p < 0.01, ***p < 0.001, ****p < 0.0001 (A-F). Data are mean ± SEM. Data are from three to four independent experiments. Each dot represents an independent experiment.

To more specifically address the role of nuclear G-actin binding to YAP, we introduced the actin-binding deficient mutations into nuclear targeted YAP^S5A^ (44) to generate YAP^S5A^ ^L65D^ and YAP^S5A^ ^L68D^, and tested their activities. YAP^S5A^ is constitutively active and locates into the nucleus (44). Consistent with this, expression of YAP^S5A^ showed enhanced transcriptional activity, whereas expression of YAP^S5A^ ^L65D^ or YAP^S5A^ ^L68D^ in which nuclear G-actin-YAP interaction was disrupted, showed impeded YAP transcriptional activity (Fig. 5c). As a negative control, the L91A mutation previously characterized as transcriptionally inactive (57), was introduced to generate YAP^S5A^ ^L91A^ (Fig. 5c). To confirm the role of nuclear G-actin-YAP interaction on YAP transcriptional activity, we generated NLS-YAP, a YAP construct fused to nuclear localization signal sequence (NLS), as well as its actin-binding deficient derivatives, NLS-YAP^L65D^, NLS-YAP^L68D^, NLS-YAP^W199Y^ for further analysis. We also generated a triple mutant NLS-YAP^DDY^, containing mutations at all three key actin-binding residues. In line with previous results, NLS-YAP elevated transcription, while the actin binding deficient NLS-YAP constructs NLS-YAP^L65D^, NLS-YAP^L68D^, NLS-YAP^W199Y^ as well as NLS-YAP^DDY^ displayed reduced transcription (Fig. 5d). NLS-YAP^L91A^ served as negative control (Fig. 5d).

To exclude the possibility that the reduced transcriptional activity of NLS-YAP^DDY^ resulted from disrupted interactions with unidentified binding partners, we performed reporter gene assays in HeLa and MDA-MB-231 cells using NLS-YAP (HeLa) or NLS-EGFP-YAP (MDA-MB-231) constructs in which actin was directly fused to the YAP N-terminus (NLS-actin-YAP^WT^ and NLS-actin-YAP^DDY^) (Fig. 5e, f). As expected, expression of NLS-YAP^WT^ but not NLS-YAP^DDY^ induced transcriptional activity (Fig. 5e, f). Interestingly, expression of the nuclear targeted actin-YAP fusion NLS-actin-YAP^WT^ further enhanced YAP activity compared to NLS-YAP (Fig. 5e, f). Consistently, generation of the actin-YAP fusion NLS-actin-YAP^DDY^ restored YAP activity, whereas NLS-YAP^DDY^ did not (Fig. 5e, f). These data support the notion that nuclear G-actin/YAP interactions promote YAP transcriptional activity.

### Predicted nuclear actin binding to YAP/TEAD4 enhances DBD exposure of TEAD4

When Hippo signalling is off, YAP translocates to the nucleus and interacts with TEAD and the YAP-TEAD complex is then recruited to specific target genes to activate transcription (17). To better understand how nuclear actin-YAP interaction facilitates transcription at the molecular level we used AlphFold3 (43) to predict potential conformational changes of TEAD4 in the absence or presence of actin. We predicted and compared the structure of TEAD4/ muscle-CAT (M-CAT) DNA (Extended Data Fig 4a), YAP/TEAD/M-CAT DNA (Fig. 6a and Extended Data Fig 4b, d) and YAP/actin/TEAD/M-CAT DNA (Fig. 6b and Extended Data Fig 4c, d). As expected, in all of our predictions, TEAD4 DNA-binding domain (DBD) was capable of interacting with M-CAT DNA (Fig. 6a, b and Extended Data Fig 4a, b, c, d, Movie 1), which is consistent with a previous report (58). Of note, the AlphaFold prediction revealed a potential four-chain α-helical interface (4α) at the C-terminus of TEAD4 located above the DBD of TEAD4 (Fig. 6a, b and Extended Data Fig 4a, Movie 1). As shown in the model, only YAP/actin but not YAP alone was able to displace the 4α interface from the TEAD4-DBD, thereby enhancing the exposure of the DBD (Fig. 6a, b and Extended Data Fig 4b, c, d, Movie 1).

**Figure 6.**
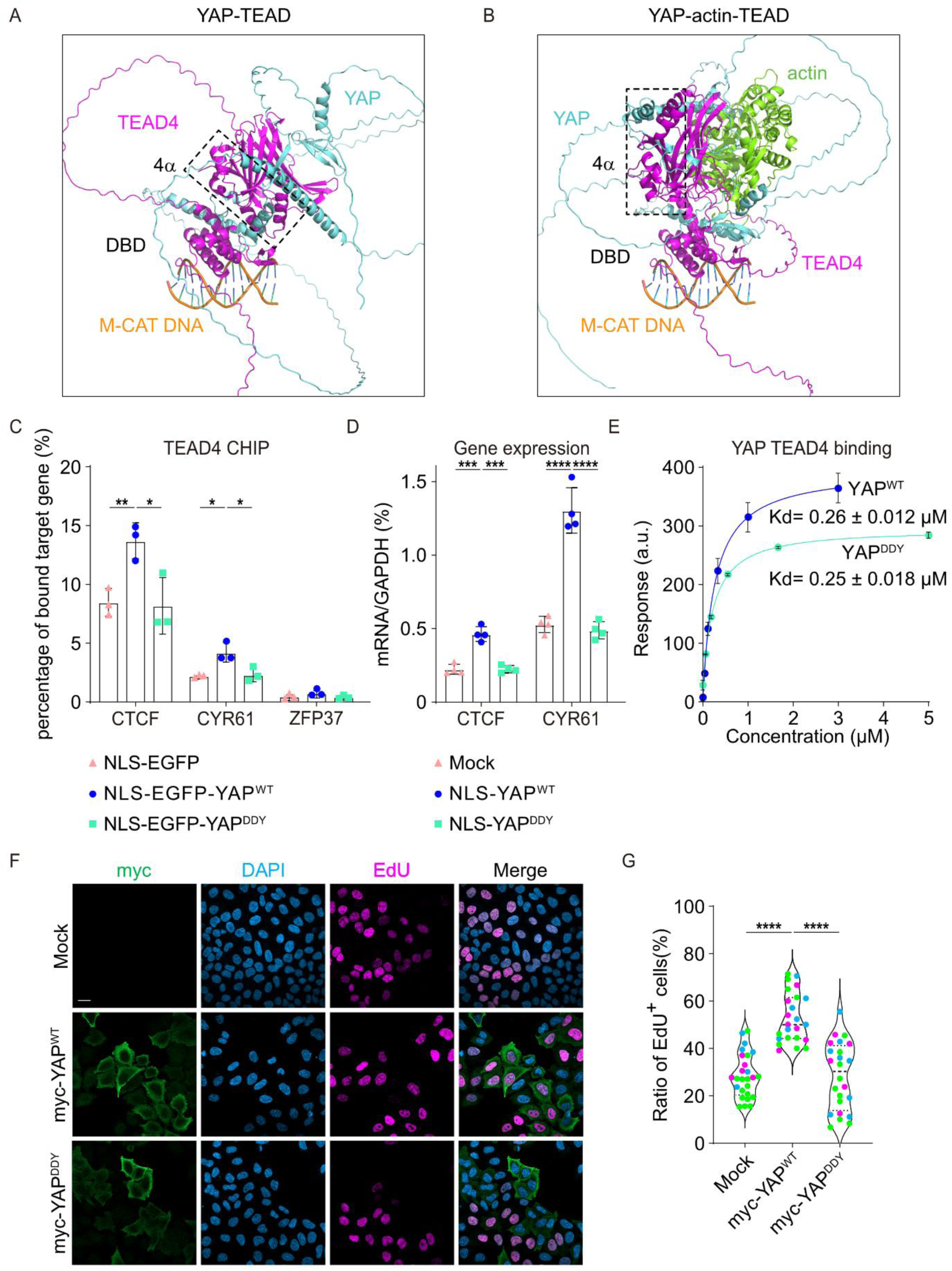
Nuclear actin-YAP binding plays a pivotal role in cell proliferation. (A-B) AlphaFold3 prediction model of TEAD4 (magenta) interaction with (A) YAP (cyan) or (B) YAP (cyan) and G-actin (green) in the presence of M-CAT DNA. The black box indicated the predicted four-chain α-helical interface (4α) interacts with TEAD4 DNA binding domain (DBD). (C) ChIP-qPCR analysis of YAP–TEAD4 recruitment to target genes in HeLa cells transfected with NLS-EGFP (pink, triangle), NLS-EGFP-YAP^WT^ (blue, circle) and NLS-EGFP-YAP^DDY^ (green, square). ZFP37 served as negative control. (D) Quantitative PCR (qPCR) analysis of YAP target gene expression in HeLa cells transfected with NLS-EGFP (pink, triangle), NLS-EGFP-YAP^WT^ (blue, circle) and NLS-EGFP-YAP^DDY^ (green, square). Data are mRNA normalized to GAPDH transcripts. Two-tailed unpaired Student’s t-test was used to calculate p-values. *p < 0.05, **p < 0.01, ***p < 0.001, ****p < 0.0001. Data are mean ± SEM. Data are from (C) three or (D) four independent experiments. Each dot represents an independent experiment. (E) Dose response curves of purified YAP^WT^ (blue) and YAP^DDY^ (green) over TEAD4-coated CM5 chip indicating the dissociation constant (K_d_). (F) Representative images of Click-iT® Plus EdU assay of HeLa cells transfected with empty plasmid (Mock), myc-YAP^WT^ and myc -YAP^DDY^ and stained for anti myc (green), DAPI (blue) and EdU (magenta). Images are shown as maximum intensity projection (MIP), Scale bar = 20 μm. (G) Violin plot with individual data points (green, magenta, cyan), median and quartiles showing the quantification of the ratio of EdU+ cells. Statistical analysis was done using One-way ANOVA with Tukeýs multiple comparison test. ****p < 0.0001. Data are from three independent experiments, n = 26, 24, 24. Each dot represents a field of view.

### Nuclear actin YAP binding regulates YAP-TEAD4 DNA binding and transcription

To test the prediction from the obtained Alphafold model, we investigated whether nuclear actin-YAP binding regulates the recruitment of YAP-TEAD to their target genes. For this we performed chromatin immunoprecipitations (ChIP) followed by qPCR in HeLa cells. Cells expressing NLS-EGFP alone were used to measure basal levels of YAP-TEAD target gene recruitment. Interestingly, we observed that expression of NLS-EGFP-YAP^WT^ significantly enhanced TEAD4 recruitment to the YAP/TEAD4 target genes CTCF and CYR61, whereas NLS-EGFP-YAP^DDY^ failed to do so (Fig. 6c). In all conditions TEAD4 recruitment to ZFP37 served as a negative control (Fig. 6c).

To further determine whether the actin-mediated modulation of DNA-binding behaviour results in a different transcriptional gene expression output, we directly analysed the common YAP/TEAD4 target genes CTCF and CYR61 by qPCR and found that NLS-YAP^WT^ but not NLS-YAP^DDY^ induced gene expression compared to the control (Fig. 6d). This was not due to the disruption of YAP-TEAD4 interaction since SPR analysis showed that the binding affinity of YAP^WT^ or YAP^DDY^ to TEAD4 was unaffected (Fig. 6e). These results demonstrate that YAP/TEAD function requires nuclear actin-YAP interaction for its DNA-binding and transcriptional activities.

### Contact inhibition is regulated by nuclear actin/YAP binding

Since high-cell density induced cell contact inhibition of proliferation is controlled by YAP/TAZ signalling(20, 22, 39, 59), we sought to determine the impact of nuclear actin/YAP binding on this process. For this, we performed Click-iT EdU cell proliferation assays on HeLa cells expressing indicated YAP constructs under confluent condition (Fig. 6f, g). The ratio of EdU+ transfected cells versus total transfected cells was used to determine the level of cell proliferation. Compared to mock transfected cells YAP significantly reduced contact inhibition, with the average EdU positive cell ratio increasing from 28% (Mock) to 53% (YAP^WT^) (Fig. 6f, g), whereas YAP^DDY^ did not alter contact inhibition with an average EdU positive cell ratio 28% (Mock) and 28% (YAP^DDY^) (Fig. 6f, g). This indicates that actin/YAP interactions are essential for control of contact inhibition versus proliferation.

## DISCUSSION

While it is well established that the cytoskeleton can regulate YAP transcriptional activity (21, 55, 60), a direct role of nuclear actin in this process has been elusive. Actin inside the nuclear compartment has been shown to be involved in transcription via its association with the RNA polymerase 2 complex (61–64), DNA repair (1, 65, 66) or via chromatin modulation (40, 67–69). In this study, we identify a direct interaction between YAP and G-actin within the cell nucleus, where YAP binds with TEAD to activate gene expression (70). We uncover that direct binding of nuclear G-actin to YAP plays a crucial role in YAP transcriptional activity. Nuclear actin can play a role in general RNA polymerase 2-mediated transcription and increased expression of nuclear actin may lead to a global enhancement of transcription (51, 62, 63). However, actin-binding deficient YAP failed to activate transcription in both, reporter gene assays and target gene qPCR analysis, strongly arguing for a specific requirement of actin for the activity of YAP.

A previously reported structure of YAP/TEAD4 (PDB: 3JUA) (71) and TEAD4/M-CAT DNA (PDB: 5GZB) (58) did not utilize full-length TEAD4. Hence, how YAP/TEAD4 interaction alters the TEAD4/DNA conformation remains at present unclear. Our predicted model using a full-length TEAD4 indicates an intramolecular interaction in TEAD4 involving its DBD and a 4α interface (Fig. 6a, b and Extended Data Fig. 4, Movie 1) while a actin/YAP complex with TEAD4 could mobilize the 4α interface to facilitate better exposure of the DBD (Fig. 6a, b and Extended Data Fig. 4, Movie 1). This scenario is in agreement with our CHIP-qPCR results showing that NLS-EGFP-YAP^WT^ but not NLS-EGFP-YAP^DDY^ enhanced the recruitment of TEAD4 to target genes (Fig. 6c). Thus, based on our findings and the AlphaFold predictions, we propose a model by which G-actin binding to YAP regulates YAP activity via two complementary mechanisms. Firstly, YAP/actin binding facilitates YAP nuclear localization. Secondly, the direct binding between G-actin and YAP is essential for promoting YAP/TEAD interaction with DNA and transcription. However, the exact structural basis underlying this mechanism remains to be elucidated.

Nuclear actin levels are implicated in various physiologically relevant processes, including stress responses, cell differentiation, and transcription (72–78). Maintaining appropriate nuclear actin levels is essential for transcription, and perturbations can alter transcriptional activity (72). At the global transcription level, elevated nuclear actin levels correlate to enhanced transcriptional activity and DNA synthesis in human S1 cells, mouse ScP2 cells and 3T3-L1 cells (73–75). In contrast, at the gene-specific level, higher nuclear actin levels exhibit different effects on gene expression, for instance, in HeLa cells and HaCaT cells, increased levels of nuclear actin downregulate MYL9, ITGB1, PAK1, while in HaCaT cells they upregulate BCL2 and NFATC1 (76, 78). Recent studies have shown that nuclear actin levels correlate with tumorigenesis. Epithelial-mesenchymal transition (EMT), a process associated with tumor initiation, invasion and metastasis, as well as EMT-associated chemotherapy resistance are both regulated by nuclear actin (79, 80). The elevated nuclear actin levels correlate with adverse prognosis (76) while nuclear actin bundling and decreased nuclear actin levels lead to cell quiescence (73, 74, 81). Moreover, the tumor repressors p53 and RASSF 1A have been suggested to regulate nuclear actin dynamics, with p53 inhibiting nuclear actin polymerization (82) and RASSF 1A suppressing nuclear actin levels (83). In addition, due to the close association between YAP and tumor progression, YAP and its transcriptional complexes have been explored as potential therapeutic targets. Verteporfin (NCT04590664) and IAG933 (NCT04857372), two YAP-TEAD interaction inhibitors, and ION537 (NCT04659096)(84), an antisense oligonucleotide targeting YAP mRNA are already in clinical trials. Taken together, our work elucidates a novel functional interaction between nuclear G-actin and YAP, that could represent a promising new target for pharmacological intervention.

## ACKNOWLEDGEMENTS

We thank laboratory members for helpful discussions. We thank Otilia Wunderlich for technical assistance.

## AUTHOR CONTRIBUTIONS

Hong Wang (Conceptualization, Investigation, Formal analysis, Methodology, Validation, Visualization, Writing – original draft, Writing – review & editing), Irma M. Jayawardana (Formal analysis, Investigation, Visualization, Writing – review & editing), Justus Fleisch (Formal analysis, Investigation, Visualization), Dennis Frank (Investigation, Formal analysis, Visualization, Writing – review & editing), Bing Zhao (Conceptualization, Investigation), Christos Kamaras (Investigation, Visualization, Writing – review & editing), Peter Gebhardt (Investigation, Visualization), Julian Knerr (Writing – review & editing), Shamphavi Sivabalasarma (Investigation), Sonja-Verena Albers (Supervision), and Robert Grosse (Conceptualization, Funding acquisition, Supervision, Resources, Methodology, Writing – original draft, Writing – review & editing).

## SUPPLEMENTARY DATA

Supplementary Data are available at NAR online.

## DATA AVAILABILITY

All the original data are provided as the supplementary files.

## Extended Data

**Extended Data Figure S1.**
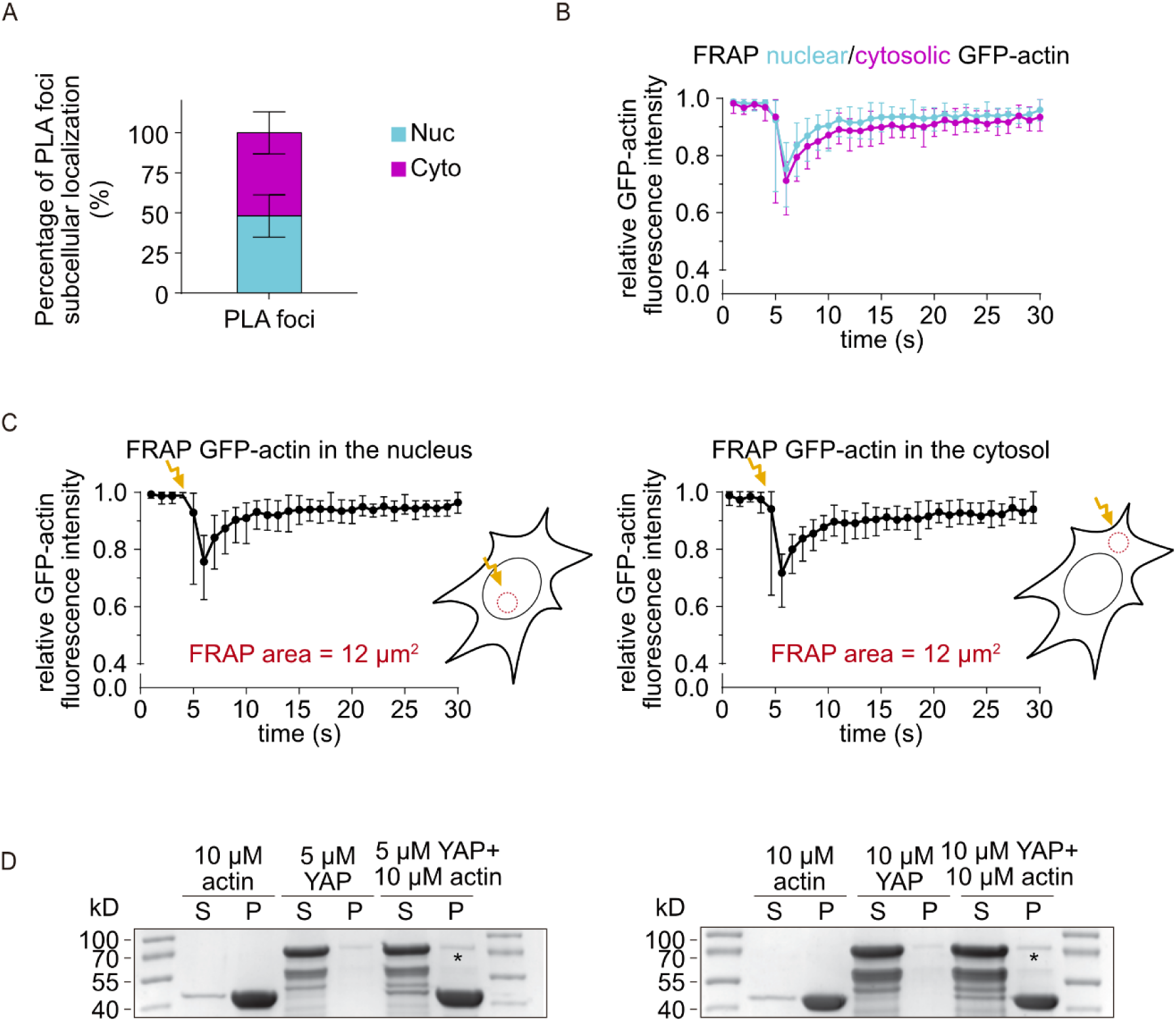
YAP binds to G-actin but not F-actin. (A)Quantification of the subcellar distribution of PLA foci. Data is n = 30 cells over three independent biological replicates. (B-C) Measurement of GFP intensity of the region of interest of FRAP. Red circle indicates photobleached region. In (B), cyan represents FRAP in nuclear compartment, and magenta represents FRAP in cytosolic compartment. Data is n = 10 cells over three independent biological replicates. (D) SDS-PAGE gels showing the supernatant (S) and pellet (P) fractions of cosedimentation assays containing 5 (left) or 10 µM (right) of the purified YAP protein and 10 µM of F-actin. Two molecular weight scale (Mw) lanes are on the both sides of the gels. These experiments were repeated three times independently with the same results.

**Extended Data Figure S2.**
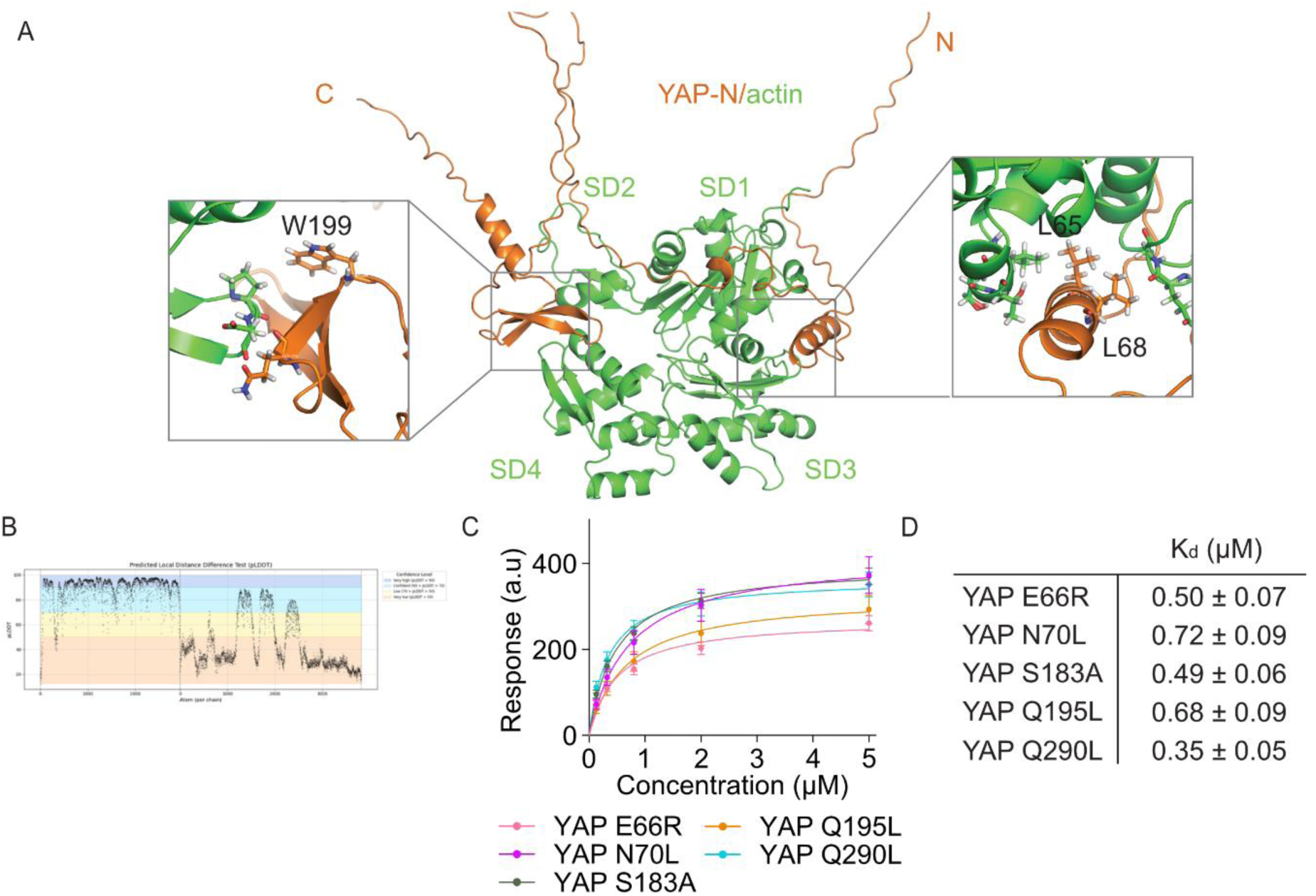
Actin binding affinity of YAP mutants. (A-B) AlphaFold 2-multimer prediction model of YAP-N (orange) interaction with G-actin (green) and (B) plddt plot. Overview (middle) and two zoom views (black box) of predicted binding interface (left and right). Predicted binding sites on YAP-N are indicated. (C-D) (C) Dose-response curves for purified indicated YAP mutants and (D) indicated dissociation constant (Kd). Data are mean ± SEM. Data are from three independent biological experiments.

**Extended Data Figure S3.**
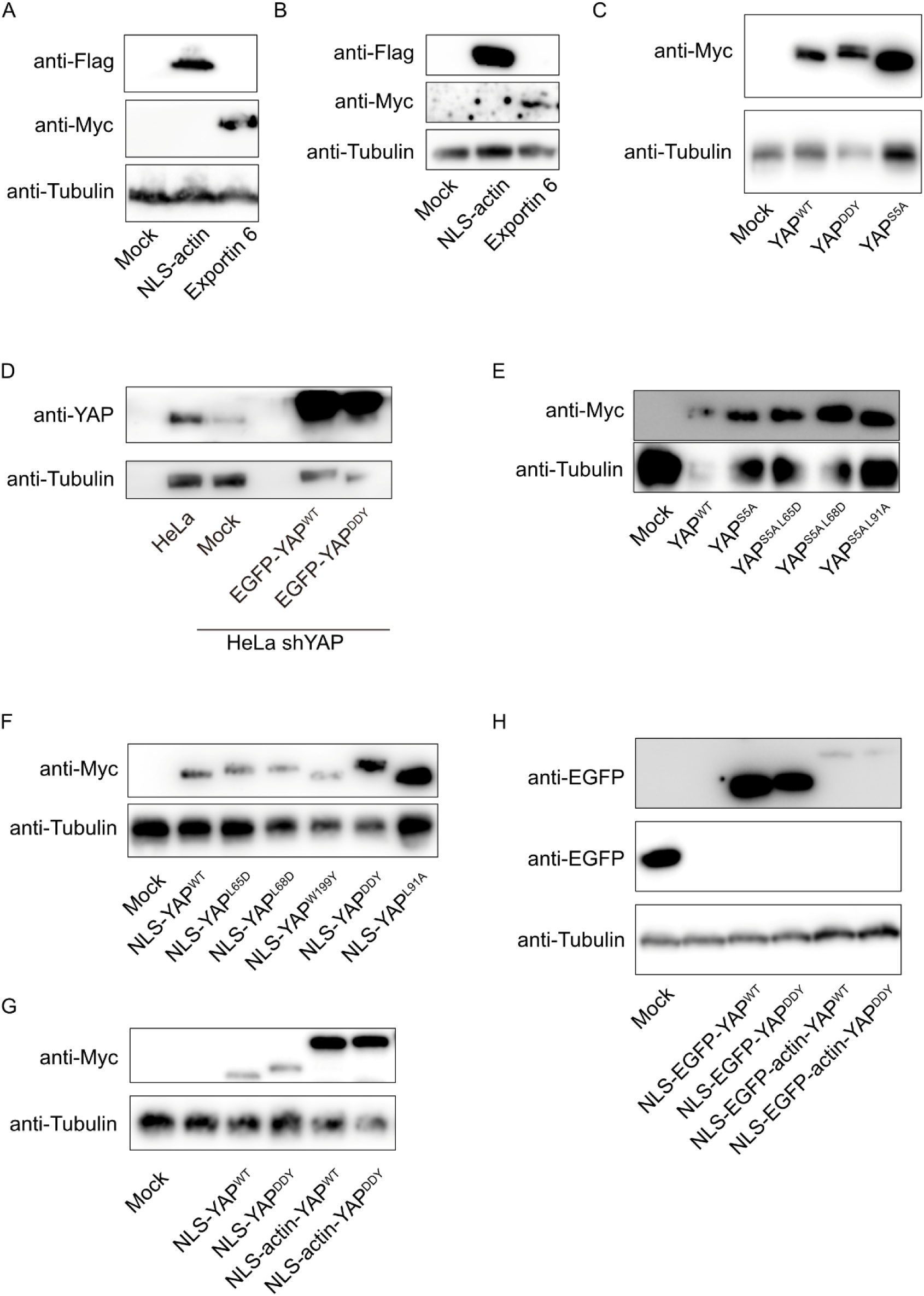
Western blot of luciferase assay. (A-H) Western blot corresponding to (A) Fig. 4C-D, (B) Fig. 4E, (C) Fig. 5A, (D) Fig. 5B, (E) Fig. 5C, (F) Fig. 5D, (G) Fig. 5E, (H) Fig. 5F.

**Extended Data Figure S4.**
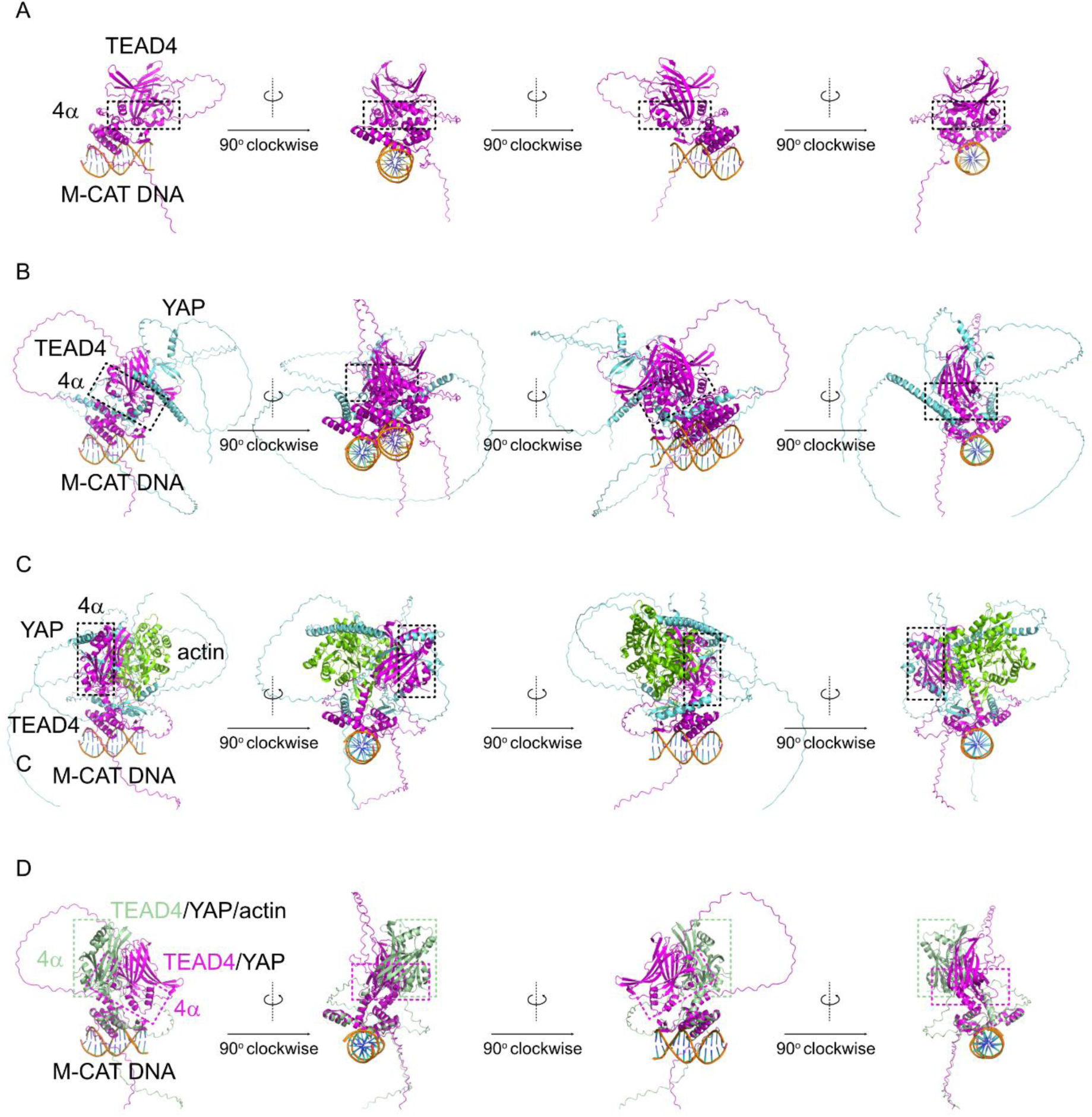
AlphaFold 3 prediction (A-C) AlphaFold 3 predicted structure of (A) TEAD4/M-CAT DNA, (B) YAP/TEAD4/M-CAT DNA and (C) YAP/G-actin/TEAD4/M-CAT DNA. TEAD4 (magenta) YAP (cyan) and actin (green). Black box indicates the 4α interface. (D) The alignment of TEAD4/M-CAT DNA from (B) YAP/TEAD4/M-CAT DNA and (C) YAP/G-actin/TEAD4/M-CAT DNA. The structure of TEAD4 and its 4α interface in the absence and presence of actin are colored in magenta and green respectively. YAP and actin are not shown.

## REFERENCES

1. Wollscheid, H.P. and Ulrich, H.D. (2023) Chromatin meets the cytoskeleton: the importance of nuclear actin dynamics and associated motors for genome stability. DNA Repair (Amst), 131, 103571.

2. Szabó, A., Borkúti, P., Kovács, Z., Kristó, I. and Vilmos, P. (2025) Recent advances in nuclear actin research. Nucleus, 16, 2498643.

3. Ulferts, S., Lopes, M., Miyamoto, K. and Grosse, R. (2024) Nuclear actin dynamics and functions at a glance. 10.1242/jcs.261630.

4. Ulferts, S., Prajapati, B., Grosse, R. and Vartiainen, M.K. (2020) Emerging Properties and Functions of Actin and Actin Filaments Inside the Nucleus. Cold Spring Harb Perspect Biol, 10.1101/cshperspect.a040121.

5. Hu, X., Liu, Z.Z., Chen, X., Schulz, V.P., Kumar, A., Hartman, A.A., Weinstein, J., Johnston, J.F., Rodriguez, E.C., Eastman, A.E., et al. (2019) MKL1-actin pathway restricts chromatin accessibility and prevents mature pluripotency activation. Nature Communications 2019 10:1, 10, 1–13.

6. Le, H.Q., Ghatak, S., Yeung, C.Y.C., Tellkamp, F., Günschmann, C., Dieterich, C., Yeroslaviz, A., Habermann, B., Pombo, A., Niessen, C.M., et al. (2016) Mechanical regulation of transcription controls Polycomb-mediated gene silencing during lineage commitment. Nature Cell Biology 2016 18:8, 18, 864–875.

7. Plessner, M. and Grosse, R. (2019) Dynamizing nuclear actin filaments. Curr Opin Cell Biol, 56, 1–6.

8. Cen, B., Selvaraj, A., Burgess, R.C., Hitzler, J.K., Ma, Z., Morris, S.W. and Prywes, R. (2003) Megakaryoblastic Leukemia 1, a Potent Transcriptional Coactivator for Serum Response Factor (SRF), Is Required for Serum Induction of SRF Target Genes. Mol Cell Biol, 23, 6597–6608.

9. Mouilleron, S., Guettler, S., Langer, C.A., Treisman, R. and McDonald, N.Q. (2008) Molecular basis for G-actin binding to RPEL motifs from the serum response factor coactivator MAL. EMBO Journal, 27, 3198–3208.

10. Mouilleron, S., Langer, C.A., Guettler, S., McDonald, N.Q. and Treisman, R. (2011) Structure of a pentavalent G-actin•MRTF-A complex reveals how G-actin controls nucleocytoplasmic shuttling of a transcriptional coactivator. Sci Signal, 4, 1–10.

11. Baarlink, C., Wang, H. and Grosse, R. (2013) Nuclear actin network assembly by formins regulates the SRF coactivator MAL. Science (1979), 340, 864–867.

12. Posern, G., Miralles, F., Guettler, S. and Treisman, R. (2004) Mutant actins that stabilise F-actin use distinct mechanisms to activate the SRF coactivator MAL. EMBO Journal, 23, 3973–3983.

13. Miralles, F., Posern, G., Zaromytidou, A.-I. and Treisman, R. (2003) Actin Dynamics Control SRF Activity by Regulation of Its Coactivator MAL. Cell, 113, 329–342.

14. Vartiainen, M.K., Guettler, S., Larijani, B. and Treisman, R. (2007) Nuclear Actin Regulates Dynamic Subcellular Localization and Activity of the SRF Cofactor MAL. Science (1979), 316, 1749–1752.

15. Olson, E.N. and Nordheim, A. (2010) Linking actin dynamics and gene transcription to drive cellular motile functions. Nature Reviews Molecular Cell Biology 2010 11:5, 11, 353–365.

16. Foster, C.T., Gualdrini, F. and Treisman, R. (2017) Mutual dependence of the MRTF-SRF and YAP-TEAD pathways in cancer-associated fibroblasts is indirect and mediated by cytoskeletal dynamics. Genes Dev, 31, 2361–2375.

17. Chen, L., Loh, P.G. and Song, H. (2010) Structural and functional insights into the TEAD-YAP complex in the Hippo signaling pathway. Protein Cell, 1, 1073–1083.

18. Heng, B.C., Zhang, X., Aubel, D., Bai, Y., Li, X., Wei, Y., Fussenegger, M. and Deng, X. (2021) An overview of signaling pathways regulating YAP/TAZ activity. Cellular and Molecular Life Sciences, 78, 497–512.

19. Fu, M., Hu, Y., Lan, T., Guan, K.L., Luo, T. and Luo, M. (2022) The Hippo signalling pathway and its implications in human health and diseases. Signal Transduction and Targeted Therapy 2022 7:1, 7, 1–20.

20. Zhao, B., Wei, X., Li, W., Udan, R.S., Yang, Q., Kim, J., Xie, J., Ikenoue, T., Yu, J., Li, L., et al. (2007) Inactivation of YAP oncoprotein by the Hippo pathway is involved in cell contact inhibition and tissue growth control. Genes Dev, 21, 2747–2761.

21. Dupont, S., Morsut, L., Aragona, M., Enzo, E., Giulitti, S., Cordenonsi, M., Zanconato, F., Le Digabel, J., Forcato, M., Bicciato, S., et al. (2011) Role of YAP/TAZ in mechanotransduction. Nature, 474, 179–184.

22. Aragona, M., Panciera, T., Manfrin, A., Giulitti, S., Michielin, F., Elvassore, N., Dupont, S. and Piccolo, S. (2013) A mechanical checkpoint controls multicellular growth through YAP/TAZ regulation by actin-processing factors. Cell, 154, 1047–1059.

23. Wang, L., Luo, J.Y., Li, B., Tian, X.Y., Chen, L.J., Huang, Y., Liu, J., Deng, D., Lau, C.W., Wan, S., et al. (2016) Integrin-YAP/TAZ-JNK cascade mediates atheroprotective effect of unidirectional shear flow. Nature 2016 540:7634, 540, 579–582.

24. Yu, F.X., Zhao, B., Panupinthu, N., Jewell, J.L., Lian, I., Wang, L.H., Zhao, J., Yuan, H., Tumaneng, K., Li, H., et al. (2012) Regulation of the Hippo-YAP pathway by G-protein-coupled receptor signaling. Cell, 150, 780–791.

25. Wada, K.I., Itoga, K., Okano, T., Yonemura, S. and Sasaki, H. (2011) Hippo pathway regulation by cell morphology and stress fibers. Development, 138, 3907–3914.

26. Chang, L., Azzolin, L., Di Biagio, D., Zanconato, F., Battilana, G., Lucon Xiccato, R., Aragona, M., Giulitti, S., Panciera, T., Gandin, A., et al. (2018) The SWI/SNF complex is a mechanoregulated inhibitor of YAP and TAZ. Nature 2018 563:7730, 563, 265–269.

27. Piccolo, S., Panciera, T., Contessotto, P. and Cordenonsi, M. (2023) YAP/TAZ as master regulators in cancer: modulation, function and therapeutic approaches. Nat Cancer, 4, 9–26.

28. Hartman, A.A., Scalf, S.M., Zhang, J., Hu, X., Chen, X., Eastman, A.E., Yang, C. and Guo, S. (2020) YAP Non-cell-autonomously Promotes Pluripotency Induction in Mouse Cells. Stem Cell Reports, 14, 730–743.

29. Khalil, A.A., Smits, D., Haughton, P.D., Koorman, T., Jansen, K.A., Verhagen, M.P., van der Net, M., van Zwieten, K., Enserink, L., Jansen, L., et al. (2024) A YAP-centered mechanotransduction loop drives collective breast cancer cell invasion. Nature Communications 2024 15:1, 15, 1–17.

30. Lamar, J.M., Stern, P., Liu, H., Schindler, J.W., Jiang, Z.G. and Hynes, R.O. (2012) The Hippo pathway target, YAP, promotes metastasis through its TEAD-interaction domain. Proc Natl Acad Sci U S A, 109, E2441–E2450.

31. He, C., Lv, X., Huang, C., Angeletti, P.C., Hua, G., Dong, J., Zhou, J., Wang, Z., Ma, B., Chen, X., et al. (2019) A Human Papillomavirus-Independent Cervical Cancer Animal Model Reveals Unconventional Mechanisms of Cervical Carcinogenesis. Cell Rep, 26, 2636–2650.e5.

32. Luo, J., Zou, H., Guo, Y., Tong, T., Chen, Y., Xiao, Y., Pan, Y. and Li, P. (2023) The oncogenic roles and clinical implications of YAP/TAZ in breast cancer. British Journal of Cancer 2023 128:9, 128, 1611–1624.

33. de Bruijn, I., Kundra, R., Mastrogiacomo, B., Tran, T.N., Sikina, L., Mazor, T., Li, X., Ochoa, A., Zhao, G., Lai, B., et al. (2023) Analysis and Visualization of Longitudinal Genomic and Clinical Data from the AACR Project GENIE Biopharma Collaborative in cBioPortal. Cancer Res, 83, 3861–3867.

34. Gao, J., Aksoy, B.A., Dogrusoz, U., Dresdner, G., Gross, B., Sumer, S.O., Sun, Y., Jacobsen, A., Sinha, R., Larsson, E., et al. (2013) Integrative analysis of complex cancer genomics and clinical profiles using the cBioPortal. Sci Signal, 6, 1–1.

35. Cerami, E., Gao, J., Dogrusoz, U., Gross, B.E., Sumer, S.O., Aksoy, B.A., Jacobsen, A., Byrne, C.J., Heuer, M.L., Larsson, E., et al. (2012) The cBio Cancer Genomics Portal: An Open Platform for Exploring Multidimensional Cancer Genomics Data. Cancer Discov, 2, 401–404.

36. Szulzewsky, F., Arora, S., Hoellerbauer, P., King, C., Nathan, E., Chan, M., Cimino, P.J., Ozawa, T., Kawauchi, D., Pajtler, K.W., et al. (2020) Comparison of tumor-associated YAP1 fusions identifies a recurrent set of functions critical for oncogenesis. Genes Dev, 34, 1051–1064.

37. Kokai, E., Beck, H., Weissbach, J., Arnold, F., Sinske, D., Sebert, U., Gaiselmann, G., Schmidt, V., Walther, P., Münch, J., et al. (2014) Analysis of nuclear actin by overexpression of wild-type and actin mutant proteins. Histochem Cell Biol, 141, 123–135.

38. Basu, S., Totty, N.F., Irwin, M.S., Sudol, M. and Downward, J. (2003) Akt Phosphorylates the Yes-Associated Protein, YAP, to Induce Interaction with 14–3-3 and Attenuation of p73-Mediated Apoptosis. Mol Cell, 11, 11–23.

39. Zhao, B., Ye, X., Yu, J., Li, L., Li, W., Li, S., Yu, J., Lin, J.D., Wang, C.-Y., Chinnaiyan, A.M., et al. (2008) TEAD mediates YAP-dependent gene induction and growth control. 10.1101/gad.1664408.

40. Baarlink, C., Plessner, M., Sherrard, A., Morita, K., Misu, S., Virant, D., Kleinschnitz, E.M., Harniman, R., Alibhai, D., Baumeister, S., et al. (2017) A transient pool of nuclear F-actin at mitotic exit controls chromatin organization. Nat Cell Biol, 19, 1389–1399.

41. Wang, H., Said, R., Nguyen-Vigouroux, C., Henriot, V., Gebhardt, P., Pernier, J., Grosse, R. and Le Clainche, C. (2024) Talin and vinculin combine their activities to trigger actin assembly. Nature Communications 2024 15:1, 15, 1–15.

42. Knerr, J., Werner, R., Schwan, C., Wang, H., Gebhardt, P., Grötsch, H., Caliebe, A., Spielmann, M., Holterhus, P.M., Grosse, R., et al. (2023) Formin-mediated nuclear actin at androgen receptors promotes transcription. Nature, 617, 616–622.

43. Abramson, J., Adler, J., Dunger, J., Evans, R., Green, T., Pritzel, A., Ronneberger, O., Willmore, L., Ballard, A.J., Bambrick, J., et al. (2024) Accurate structure prediction of biomolecular interactions with AlphaFold 3. Nature, 630, 493–500.

44. Jumper, J., Evans, R., Pritzel, A., Green, T., Figurnov, M., Ronneberger, O., Tunyasuvunakool, K., Bates, R., Žídek, A., Potapenko, A., et al. (2021) Highly accurate protein structure prediction with AlphaFold. Nature 2021 >596:7873, 596, 583–589.

45. Evans, R., O’Neill, M., Pritzel, A., Antropova, N., Senior, A., Green, T., Žídek, A., Bates, R., Blackwell, S., Yim, J., et al. (2021) Protein complex prediction with AlphaFold-Multimer. 10.1101/2021.10.04.463034.

46. Mirdita, M., Schütze, K., Moriwaki, Y., Heo, L., Ovchinnikov, S. and Steinegger, M. (2022) ColabFold: making protein folding accessible to all. Nat Methods, 19, 679–682.

47. Tosic, J., Kim, G.J., Pavlovic, M., Schröder, C.M., Mersiowsky, S.L., Barg, M., Hofherr, A., Probst, S., Köttgen, M., Hein, L., et al. (2019) Eomes and Brachyury control pluripotency exit and germ-layer segregation by changing the chromatin state. Nature Cell Biology 2019 21:12, 21, 1518– 1531.

48. Schröder, C.M., Zissel, L., Mersiowsky, S.L., Tekman, M., Probst, S., Schüle, K.M., Preissl, S., Schilling, O., Timmers, H.T.M. and Arnold, S.J. (2025) EOMES establishes mesoderm and endoderm differentiation potential through SWI/SNF-mediated global enhancer remodeling. Dev Cell, 60, 735–748.e5.

49. Stein, C., Bardet, A.F., Roma, G., Bergling, S., Clay, I., Ruchti, A., Agarinis, C., Schmelzle, T., Bouwmeester, T., Schübeler, D., et al. (2015) YAP1 Exerts Its Transcriptional Control via TEAD-Mediated Activation of Enhancers. PLoS Genet, 11, e1005465.

50. Hegazy, M., Cohen-Barak, E., Koetsier, J.L., Najor, N.A., Arvanitis, C., Sprecher, E., Green, K.J. and Godsel, L.M. (2020) Proximity Ligation Assay for Detecting Protein-Protein Interactions and Protein Modifications in Cells and Tissues in Situ. Curr Protoc Cell Biol, 89, e115.

51. Svenja Ulferts and Robert Grosse (2024) SUN2 mediates calcium-triggered nuclear actin polymerization to cluster active RNA polymerase II. bioRxiv.

52. Guettler, S., Vartiainen, M.K., Miralles, F., Larijani, B. and Treisman, R. (2008) RPEL Motifs Link the Serum Response Factor Cofactor MAL but Not Myocardin to Rho Signaling via Actin Binding. Mol Cell Biol, 28, 732–742.

53. Sudol, M., Shields, D.C. and Farooq, A. (2012) Structures of YAP protein domains reveal promising targets for development of new cancer drugs. Semin Cell Dev Biol, 23, 827–833.

54. Vartiainen, M.K., Guettler, S., Larijani, B. and Treisman, R. (2007) Nuclear actin regulates dynamic subcellular localization and activity of the SRF cofactor MAL. Science (1979), 316, 1749–1752.

55. Wada, K.I., Itoga, K., Okano, T., Yonemura, S. and Sasaki, H. (2011) Hippo pathway regulation by cell morphology and stress fibers. Development, 138, 3907–3914.

56. Stüven, T., Hartmann, E. and Görlich, D. (2003) Exportin 6: a novel nuclear export receptor that is specific for profilin·actin complexes. EMBO J, 22, 5928–5940.

57. Li, Z., Zhao, B., Wang, P., Chen, F., Dong, Z., Yang, H., Guan, K.L. and Xu, Y. (2010) Structural insights into the YAP and TEAD complex. Genes Dev, 24, 235–240.

58. Shi, Z., He, F., Chen, M., Hua, L., Wang, W., Jiao, S. and Zhou, Z. (2017) DNA-binding mechanism of the Hippo pathway transcription factor TEAD4. Oncogene 2017 36:30, 36, 4362–4369.

59. Pavel, M., Renna, M., Park, S.J., Menzies, F.M., Ricketts, T., Füllgrabe, J., Ashkenazi, A., Frake, R.A., Lombarte, A.C., Bento, C.F., et al. (2018) Contact inhibition controls cell survival and proliferation via YAP/TAZ-autophagy axis. Nature Communications 2018 9:1, 9, 1–18.

60. Zhao, B., Li, L., Wang, L., Wang, C.-Y., Yu, J. and Guan, K.-L. (2012) Cell detachment activates the Hippo pathway via cytoskeleton reorganization to induce anoikis. Genes Dev, 26, 54–68.

61. Hu, P., Wu, S. and Hernandez, N. (2004) A role for β-actin in RNA polymerase III transcription. Genes Dev, 18, 3010–3015.

62. Wei, M., Fan, X., Ding, M., Li, R., Shao, S., Hou, Y., Meng, S., Tang, F., Li, C. and Sun, Y. (2020) Nuclear actin regulates inducible transcription by enhancing RNA polymerase II clustering. Sci Adv, 6.

63. Hofmann, W.A., Stojiljkovic, L., Fuchsova, B., Vargas, G.M., Mavrommatis, E., Philimonenko, V., Kysela, K., Goodrich, J.A., Lessard, J.L., Hope, T.J., et al. (2004) Actin is part of pre-initiation complexes and is necessary for transcription by RNA polymerase II. Nat Cell Biol, 6, 1094–1101.

64. Philimonenko, V. V., Zhao, J., Iben, S., Dingová, H., Kyselá, K., Kahle, M., Zentgraf, H., Hofmann, W.A., de Lanerolle, P., Hozák, P., et al. (2004) Nuclear actin and myosin I are required for RNA polymerase I transcription. Nat Cell Biol, 6, 1165–1172.

65. Schrank, B.R., Aparicio, T., Li, Y., Chang, W., Chait, B.T., Gundersen, G.G., Gottesman, M.E. and Gautier, J. (2018) Nuclear ARP2/3 drives DNA break clustering for homology-directed repair. Nature 2018 559:7712, 559, 61–66.

66. Caridi, C.P., D’agostino, C., Ryu, T., Zapotoczny, G., Delabaere, L., Li, X., Khodaverdian, V.Y., Amaral, N., Lin, E., Rau, A.R., et al. (2018) Nuclear F-actin and myosins drive relocalization of heterochromatic breaks. Nature 2018 559:7712, 559, 54–60.

67. Zhang, X., Wang, X., Zhang, Z. and Cai, G. (2019) Structure and functional interactions of INO80 actin/Arp module. J Mol Cell Biol, 11, 345–355.

68. Nishimoto, N., Watanabe, M., Watanabe, S., Sugimoto, N., Yugawa, T., Ikura, T., Koiwai, O., Kiyono, T. and Fujita, M. (2012) Heterocomplex Formation by Arp4 and β-Actin Involved in Integrity of the Brg1 Chromatin Remodeling Complex. J Cell Sci, 10.1242/jcs.104349.

69. Mashtalir, N., D’Avino, A.R., Michel, B.C., Luo, J., Pan, J., Otto, J.E., Zullow, H.J., McKenzie, Z.M., Kubiak, R.L., St. Pierre, R., et al. (2018) Modular Organization and Assembly of SWI/SNF Family Chromatin Remodeling Complexes. Cell, 175, 1272–1288.e20.

70. Pocaterra, A., Romani, P. and Dupont, S. (2020) YAP/TAZ functions and their regulation at a glance. J Cell Sci, 133.

71. Chen, L., Chan, S.W., Zhang, X.Q., Walsh, M., Lim, C.J., Hong, W. and Song, H. (2010) Structural basis of YAP recognition by TEAD4 in the Hippo pathway. Genes Dev, 24, 290–300.

72. Hyrskyluoto, A. and Vartiainen, M.K. (2020) Regulation of nuclear actin dynamics in development and disease This review comes from a themed issue on Cell Nucleus. Curr Opin Cell Biol, 2020, 18–24.

73. Spencer, V.A., Costes, S., Inman, J.L., Xu, R., Chen, J., Hendzel, M.J. and Bissell, M.J. (2011) Depletion of nuclear actin is a key mediator of quiescence in epithelial cells. J Cell Sci, 124, 123–132.

74. Fiore, A.P.Z.P., Spencer, V.A., Mori, H., Carvalho, H.F., Bissell, M.J. and Bruni-Cardoso, A. (2017) Laminin-111 and the Level of Nuclear Actin Regulate Epithelial Quiescence via Exportin-6. Cell Rep, 19, 2102–2115.

75. Dopie, J., Skarp, K.P., Rajakylä, E.K., Tanhuanpää, K. and Vartiainen, M.K. (2012) Active maintenance of nuclear actin by importin 9 supports transcription. Proc Natl Acad Sci U S A, 109, E544–E552.

76. Chatzifrangkeskou, M., Pefani, D., Eyres, M., Vendrell, I., Fischer, R., Pankova, D. and O’Neill, E. (2019) RASSF 1A is required for the maintenance of nuclear actin levels. EMBO J, 38.

77. Huet, G., Skarp, K.P. and Vartiainen, M.K. (2012) Nuclear actin levels as an important transcriptional switch. Transcription, 3, 226–230.

78. Sharili, A.S., Kenny, F.N., Vartiainen, M.K. and Connelly, J.T. (2016) Nuclear actin modulates cell motility via transcriptional regulation of adhesive and cytoskeletal genes. Sci Rep, 6, 1–9.

79. Du, W.W., Qadir, J., Du, K.Y., Chen, Y., Wu, N. and Yang, B.B. (2023) Nuclear Actin Polymerization Regulates Cell Epithelial-Mesenchymal Transition. Advanced Science, 10, 2300425.

80. Debaugnies, M., Rodríguez-Acebes, S., Blondeau, J., Parent, M.-A., Zocco, M., Song, Y., de Maertelaer, V., Moers, V., Latil, M., Dubois, C., et al. (2023) RHOJ controls EMT-associated resistance to chemotherapy. Nature, 616, 168–175.

81. Lawson, C.D., Peel, S., Jayo, A., Corrigan, A., Iyer, P., Dalrymple, M.B., Marsh, R.J., Cox, S., Van Audenhove, I., Gettemans, J., et al. (2022) Nuclear fascin regulates cancer cell survival. Elife, 11.

82. Torii, T., Sugimoto, W., Itoh, K., Kinoshita, N., Gessho, M., Goto, T., Uehara, I., Nakajima, W., Budirahardja, Y., Miyoshi, D., et al. (2023) Loss of p53 function promotes DNA damage-induced formation of nuclear actin filaments. Cell Death Dis, 14, 766.

83. Chatzifrangkeskou, M., Pefani, D., Eyres, M., Vendrell, I., Fischer, R., Pankova, D. and O’Neill, E. (2019) RASSF 1A is required for the maintenance of nuclear actin levels. EMBO J, 38.

84. Papavassiliou, K.A., Sofianidi, A.A. and Papavassiliou, A.G. (2024) YAP/TAZ-TEAD signalling axis: A new therapeutic target in malignant pleural mesothelioma. J Cell Mol Med, 28, e18330.

