## Extended Data for "Regulation of YAP activity by nuclear G-actin binding"

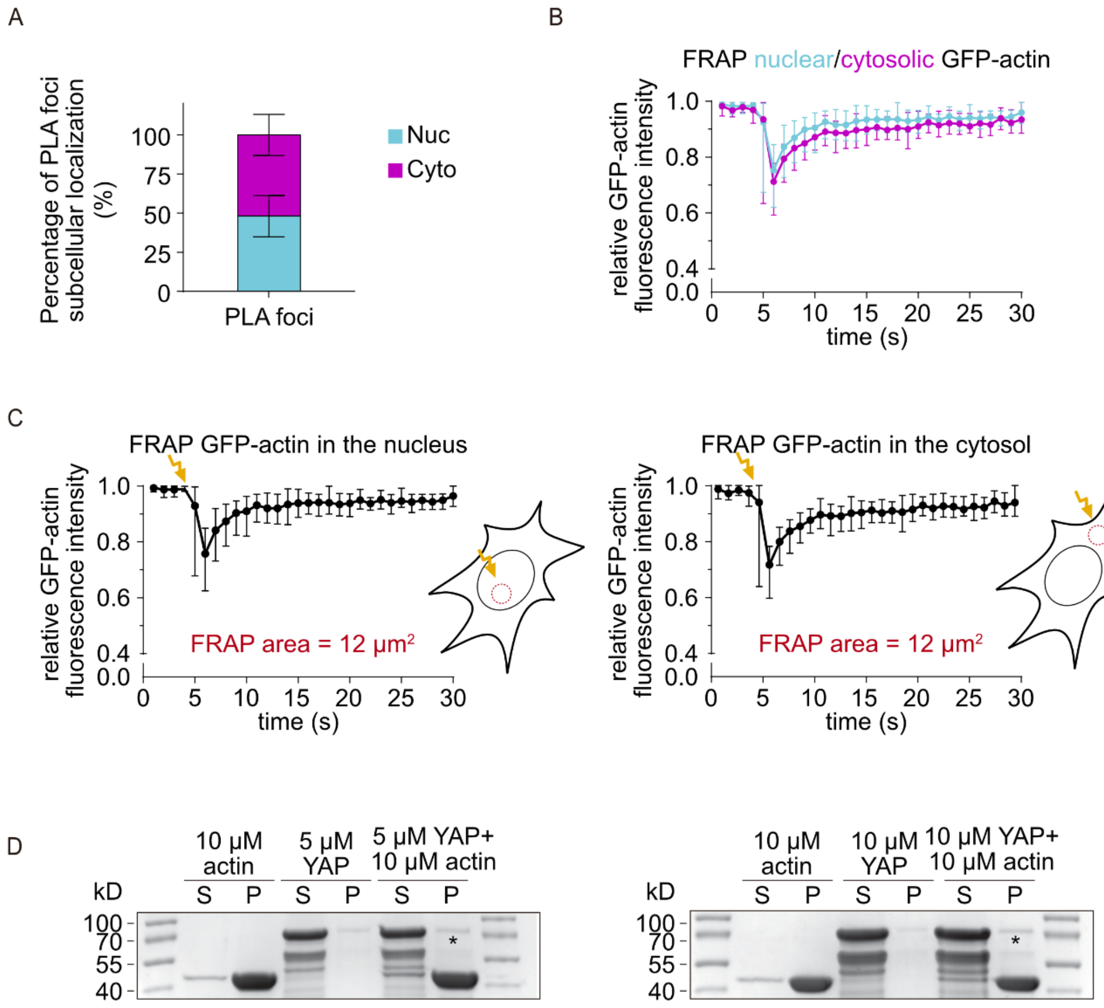


**Extended Data Figure S1.** YAP binds to G-actin but not F-actin. (A)Quantification of the subcellar distribution of PLA foci. Data is n = 30 cells over three independent biological replicates. (B-C) Measurement of GFP intensity of the region of interest of FRAP. Red circle indicates photobleached region. In (B), cyan represents FRAP in nuclear compartment, and magenta represents FRAP in cytosolic compartment. Data is n = 10 cells over three independent biological replicates. (D) SDS-PAGE gels showing the supernatant (S) and pellet (P) fractions of cosedimentation assays containing 5 (left) or 10 µM (right) of the purified YAP protein and 10 µM of F-actin. Two molecular weight scale (Mw) lanes are on the both sides of the gels. These experiments were repeated three times independently with the same results.

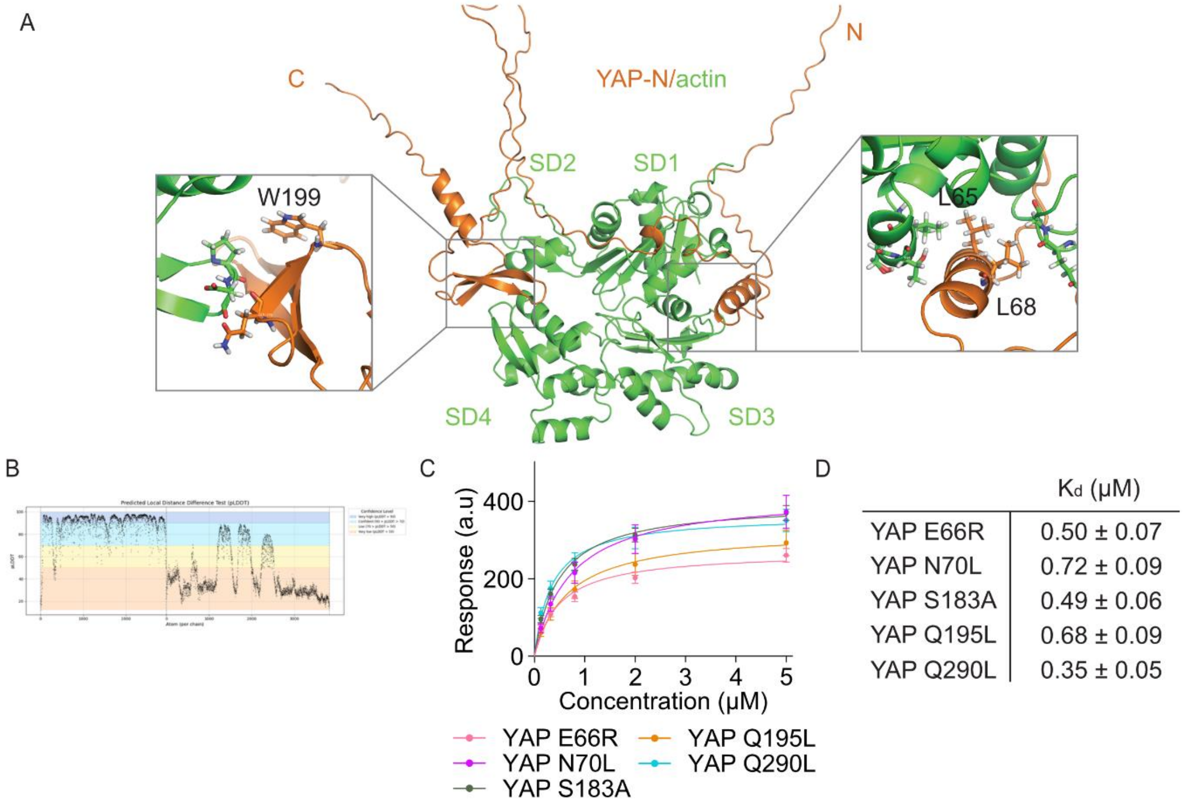


| | $K_d$ (µM) |
|---|---|
| YAP E66R | 0.50 ± 0.07 |
| YAP N70L | 0.72 ± 0.09 |
| YAP S183A | 0.49 ± 0.06 |
| YAP Q195L | 0.68 ± 0.09 |
| YAP Q290L | 0.35 ± 0.05 |

**Extended Data Figure S2.** Actin binding affinity of YAP mutants. (A-B) AlphaFold 2-multimer prediction model of YAP-N (orange) interaction with G-actin (green) and (B) plddt plot. Overview (middle) and two zoom views (black box) of predicted binding interface (left and right). Predicted binding sites on YAP-N are indicated. (C-D) (C) Dose-response curves for purified indicated YAP mutants and (D) indicated dissociation constant (Kd). Data are mean ± SEM. Data are from three independent biological experiments.

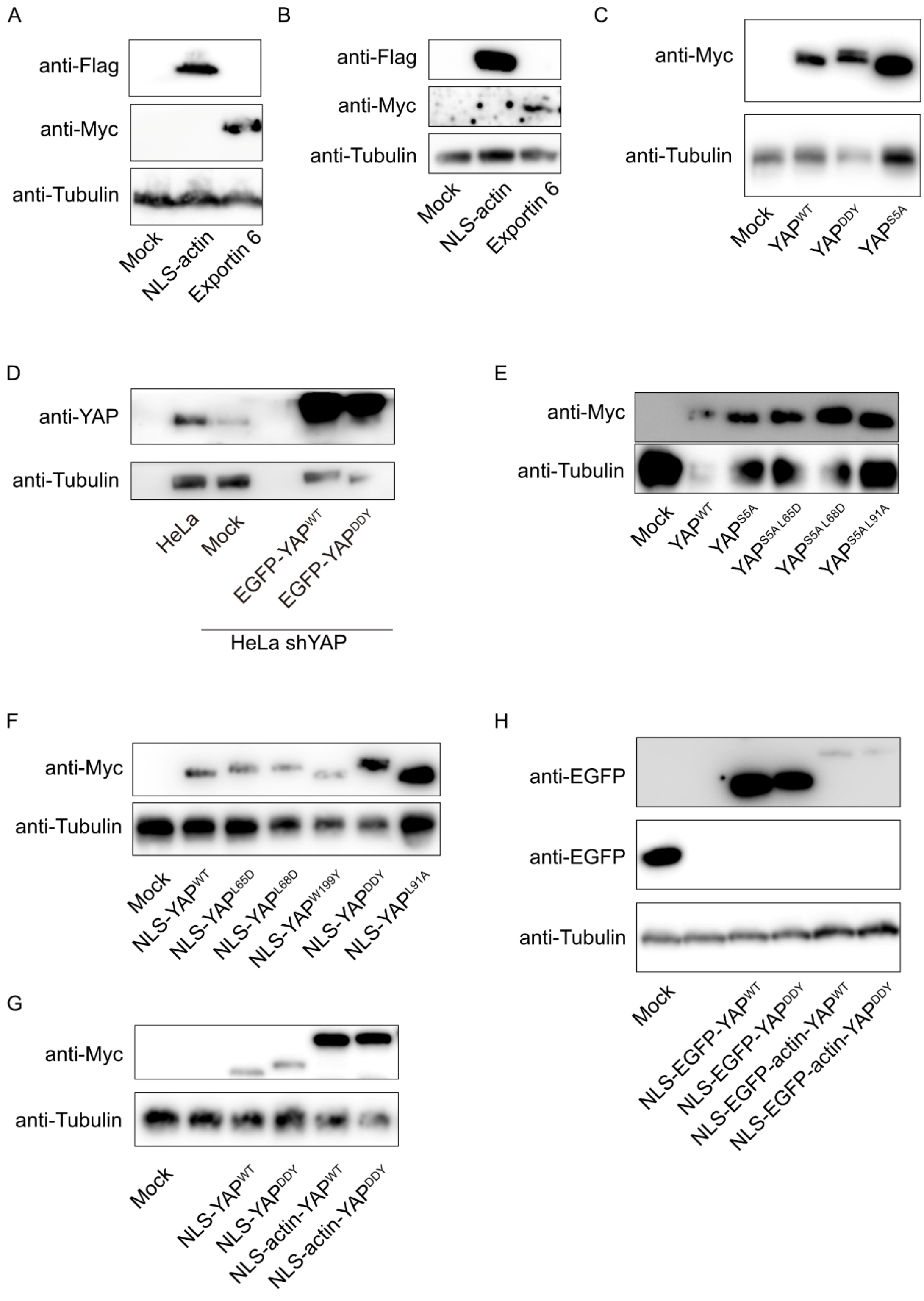


**Extended Data Figure S3.** Western blot of luciferase assay. (A-H) Western blot corresponding to (A) Fig. 4C-D, (B) Fig. 4E, (C) Fig. 5A, (D) Fig. 5B, (E) Fig. 5C, (F) Fig. 5D, (G) Fig. 5E, (H) Fig. 5F.

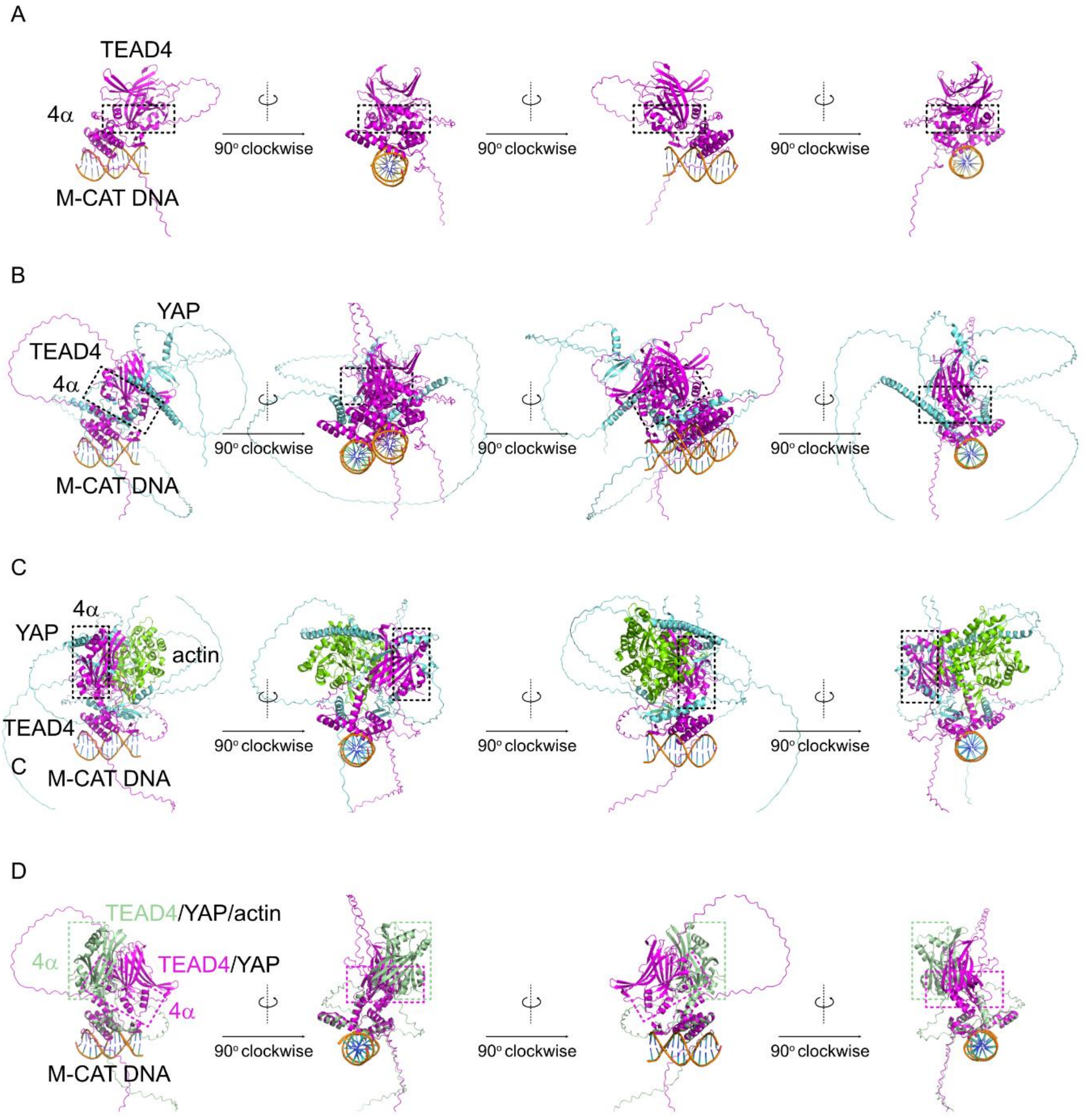


**Extended Data Figure S4.** AlphaFold 3 prediction (A-C) AlphaFold 3 predicted structure of (A) TEAD4/M-CAT DNA, (B) YAP/TEAD4/M-CAT DNA and (C) YAP/G-actin/TEAD4/M-CAT DNA. TEAD4 (magenta) YAP (cyan) and actin (green). Black box indicates the 4$\alpha$ interface. (D) The alignment of TEAD4/M-CAT DNA from (B) YAP/TEAD4/M-CAT DNA and (C) YAP/G-actin/TEAD4/M-CAT DNA. The structure of TEAD4 and its 4$\alpha$ interface in the absence and presence of actin are colored in magenta and green respectively. YAP and actin are not shown.
